# Interplay between LHCSR proteins and state transitions governs the NPQ response in intact cells of *Chlamydomonas* during light fluctuations

**DOI:** 10.1101/2021.12.31.474662

**Authors:** Collin J. Steen, Adrien Burlacot, Audrey H. Short, Krishna K. Niyogi, Graham R. Fleming

**Author notes:** Authors have had an equally valued contribution to this work (Collin J. Steen, Adrien Burlacot). **Author contributions:** C.J.S., A.B. and G.R.F. designed the research; C.J.S., A.B. and A.H.S. performed research; C.J.S., A.B., A.H.S. and G.R.F. analyzed data; C.J.S. and A.B. wrote the paper with inputs from A.H.S., K.K.N. and G.R.F.

## Abstract

Photosynthetic organisms use sunlight as the primary energy source to fix CO_2_. However, in the environment, light energy fluctuates rapidly and often exceeds saturating levels for periods ranging from seconds to hours, which can lead to detrimental effects for cells. Safe dissipation of excess light energy occurs primarily by non-photochemical quenching (NPQ) processes. In the model green microalga *Chlamydomonas reinhardtii*, photoprotective NPQ is mostly mediated by pH-sensing light-harvesting complex stress-related (LHCSR) proteins and the redistribution of light-harvesting antenna proteins between the photosystems (state transition). Although each component underlying NPQ has been documented, their relative contributions to the dynamic functioning of NPQ under fluctuating light conditions remains unknown. Here, by monitoring NPQ throughout multiple high light-dark cycles with fluctuation periods ranging from 1 to 10 minutes, we show that the dynamics of NPQ depend on the frequency of light fluctuations. Mutants impaired in the accumulation of LHCSRs (*npq4, lhcsr1*, and *npq4lhcsr1*) showed significantly less quenching during illumination, demonstrating that LHCSR proteins are responsible for the majority of NPQ during repetitive exposure to high light fluctuations. Activation of NPQ was also observed during the dark phases of light fluctuations, and this was exacerbated in mutants lacking LHCSRs. By analyzing 77K chlorophyll fluorescence spectra and chlorophyll fluorescence lifetimes and yields in a mutant impaired in state transition, we show that this phenomenon arises from state transition. Finally, we quantified the contributions of LHCSRs and state transition to the overall NPQ amplitude and dynamics for all light periods tested and show that both processes interact to facilitate NPQ during light fluctuations. We further assess the role of LHCSRs in cell growth under various periods of fluctuating light. These results highlight the dynamic functioning of photoprotection under light fluctuations and open a new way to systematically characterize the photosynthetic response to an ever-changing light environment.

**One sentence summary:** The roles of LHCSR and STT7 in NPQ vary with the light fluctuation period and duration of light fluctuation.

## Introduction

Most life on Earth is sustained by photosynthetic organisms that capture sunlight energy to convert CO_2_ and water into chemical energy. Light is captured by light-harvesting antenna complexes that contain a network of pigments absorbing photons and funneling the energy towards photosystems II and I that use it to perform photochemical reactions. Under light-limiting conditions, efficient light harvesting is crucial for maximizing the rate of CO_2_ fixation (Björkman and Demmig, 1987). However, high light (HL) intensities can saturate reaction centers and lead to the build-up of excess excitation energy, which, if unchecked, can lead to the production of reactive oxygen species and damage to both reaction centers (Khorobrykh *et al*., 2020). In nature, light exposure rapidly fluctuates in intensity with periods of HL ranging from milliseconds to hours (Graham *et al*., 2017), requiring photosynthesis to acclimate to different frequencies of HL fluctuations. For each period of HL acclimation, photosynthetic organisms exhibit photoprotective mechanisms that regulate light harvesting and safely remove excess energy (Erickson *et al*., 2015; Pinnola and Bassi, 2018; Roach, 2020).

Upon light absorption, the energy can be dissipated as heat in a process called non-photochemical quenching (NPQ). NPQ involves five components, each of which has been distinguished by its time of induction and relaxation during transition between dark and HL (Erickson et al., 2015). The fastest component, called energy-dependent quenching (qE), is triggered by luminal acidification (Briantais *et al*., 1979) and is induced and relaxed within seconds. State transition (qT) occurs within minutes and involves the phosphorylation of light-harvesting complexes (LHCs) (Allen, 1992) resulting in their detachment from Photosystem (PS) II and subsequent aggregation in a quenched state and/or association to PSI (Nagy *et al*., 2014; Ünlü *et al*., 2014; Nawrocki *et al*., 2016). Zeaxanthin-dependent quenching (qZ) requires the accumulation of zeaxanthin and probably involves quenching in the minor LHCs of PSII (Dall’Osto *et al*., 2005; Wehner *et al*., 2006; Nilkens *et al*., 2010). On longer time scales, two more sustained forms of NPQ occur: qH that takes places it the antennae of PSII (Malnoë *et al*., 2018) directly in the LHCII trimers (Bru *et al*., 2021) and photoinhibition (qI) that occurs when degradation of the PSII core protein D1 exceeds its capacity for repair due to excess excitation energy (Aro *et al*., 1993).

In the green microalga *Chlamydomonas reinhardtii*, qE is mediated by pigment-binding LHC stress-related (LHCSR) proteins (Peers *et al*., 2009; Rochaix and Bassi, 2019). LHCSRs contain protonatable residues, which sense the decreasing luminal pH generated under HL conditions (Ballottari *et al*., 2016; Tian *et al*., 2019); the protonation of LHCSRs triggers NPQ within the protein (Liguori *et al*., 2013; Kondo *et al*., 2017; Troiano *et al*., 2021), allowing fast activation of qE. While there are two types of LHCSR proteins (LHCSR1 and LHCSR3), both of which bind pigments (Bonente *et al*., 2011; Perozeni *et al*., 2020), LHCSR3 (for which two homologs are present in *Chlamydomonas*) is thought to be the main actor in qE (Peers et al., 2009; Truong, 2011). On the other hand, qT is activated by the buildup of reducing equivalents in the thylakoid membrane, which activates a serine/threonine-protein kinase (STT7) (Depege *et al*., 2003; Lemeille *et al*., 2009) that phosphorylates LHCII, enabling it to detach from PSII and ultimately reattach to PSI (Iwai *et al*., 2010a; Minagawa, 2011). While qZ has been described in *Chlamydomonas* (Niyogi *et al*., 1997), it does not seem to play a significant role in overall NPQ (Girolomoni *et al*., 2019; Tian et al., 2019), and its potential mechanism of action remains to be determined. Finally, while qH has not been described in *Chlamydomonas*, qI occurs upon continued excess illumination (Aro et al., 1993; Erickson et al., 2015) at the level of the PSII center, where oxygen-mediated sensitization creates the irreversible formation of a quenching site (Nawrocki *et al*., 2021).

The photophysical and biochemical bases for NPQ have been studied for decades (Erickson et al., 2015), however the *in vivo* operation has mostly been assessed under a single dark-to-HL transition (Nedbal and Lazár, 2021) leaving our understanding of photoprotection under more complex light patterns limited. While LHCSR and STT7 activity are both known to be important for steady-state NPQ based on measurements performed under non-repeating HL/dark periods (Allorent *et al*., 2013), their relative contributions to NPQ have not been quantified, and their response to faster-timescale fluctuating light remains unstudied. Recent work has started looking at the response of NPQ to some specific light fluctuations in *Chlamydomonas* (Roach, 2020) and in the moss *Physcomitrella* (Gao *et al*., 2021). However, the physiological role and the functioning of the NPQ components under the wide diversity of light patterns that are present in the natural environment is unexplored.

Here we utilized two distinct methods to monitor chlorophyll fluorescence in intact cells of *Chlamydomonas* that were exposed to varying frequencies of fluctuating light with HL/dark periods ranging from 1 to 10 min (**Fig. 1**). The roles of qE and qT were investigated using single or double mutants of LHCSRs and STT7. Our analysis of LHCSR mutants (*npq4 (Peers et al*., *2009), lhcsr1* (Truong, 2011), and *npq4lhcsr1* (Truong, 2011)) revealed that LHCSR3 is the main contributor to the NPQ response during the HL phase of light fluctuations. Using mutants impaired in state transition (*stt7-9 (Depege et al*., *2003)* and *stt7npq4 (Allorent et al*., *2013)*), we showed that qT quenching occurs primarily during the dark portion of the fluctuating light sequence and represents a significant part of NPQ during repeated light fluctuations. Our results showed that while qE and qT sustain most of the NPQ throughout light fluctuations, their relative importance varies during different phases of the fluctuating light response, with qT playing a larger role during dark periods and after repeated HL-dark fluctuations. Surprisingly, the light fluctuation period did not seem to have a major impact on the respective contributions of qE and qT although the contribution of qE during the dark phase was period dependent. Nonetheless, the various components of NPQ are not completely independent, and we reveal an interplay between LHCSR- and STT7-mediated NPQ that enables the wild-type photoprotective response. We further show that while *stt7* mutants are not impaired in growth under light fluctuations, short time scale light fluctuations highly impair LHCSR mutants. These findings represent an important first step in investigating the photosynthetic response to the diversity of HL periods that occur in nature.

**Figure 1.**
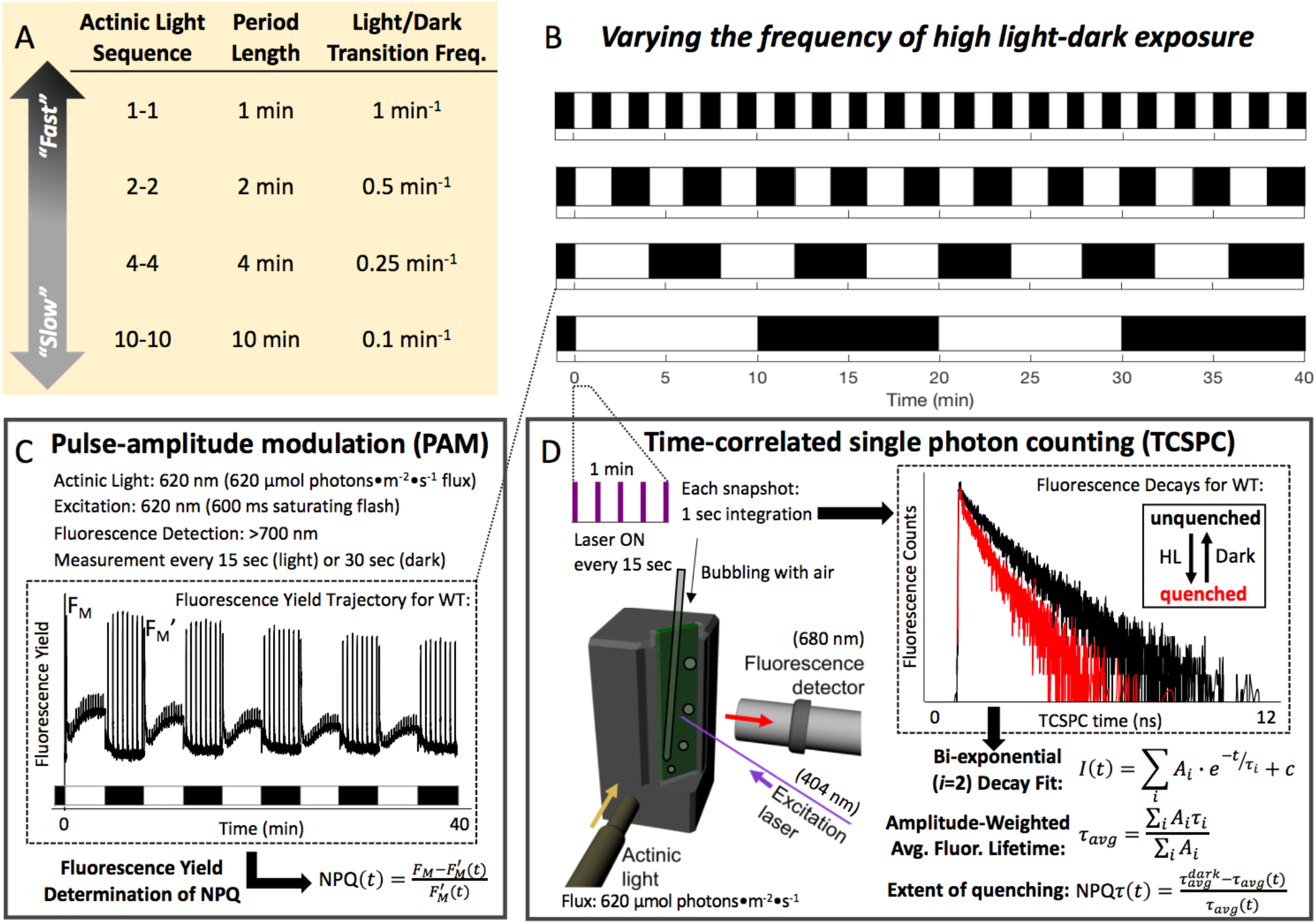
Experimental design for Chl *a* fluorescence measurements throughout exposure of *Chlamydomonas* cells to fluctuating light with various periods of HL-dark exposure. (**A, B**) Representation of the HL-dark cycles used for the 40 minutes of light fluctuation and their corresponding period, frequency, and name used throughout the main text. NPQ was measured using Pulsed Amplitude Modulation (PAM, **C**) and Time-Correlated Single Photon Counting (TCSPC, **D**). (**C**) Characteristics of the PAM measurement and representative data of fluorescence yield in WT cells. (**D**) Characteristics of the TCSPC apparatus. Shown are two decays representative of two snapshots taken in a quenched and unquenched state.

## Results

### Varying light fluctuation periods affect the dynamic NPQ response

While the photosynthetic response of *Chlamydomonas* to some light fluctuations has been reported (Cantrell and Peers, 2017; Roach, 2020), an analysis of NPQ for various periods of light fluctuations is lacking. We therefore measured chlorophyll fluorescence during light-dark cycles of 40 min with individual fluctuation periods ranging from 1 min to 10 min (**Fig. 1**). Chlorophyll fluorescence yield was measured using pulse-amplitude modulated (PAM) fluorometry and used to calculate NPQ (Klughammer and Schreiber, 2008). In tandem experiments, time-correlated single photon counting (TCSPC) was used to measure the chlorophyll fluorescence lifetime (Amarnath *et al*., 2012), which was used to calculate NPQτ (Sylak-Glassman *et al*., 2014). For all periods of light fluctuations in the wild-type strain, NPQ quickly turned on upon illumination but turned off more slowly (**Fig. 2**). The same trend was observed in NPQτ (**Fig. 2**). The 1 min period light fluctuation led to a nearly square-like response of NPQ and NPQτ (**Fig. 2**).

**Figure 2.**
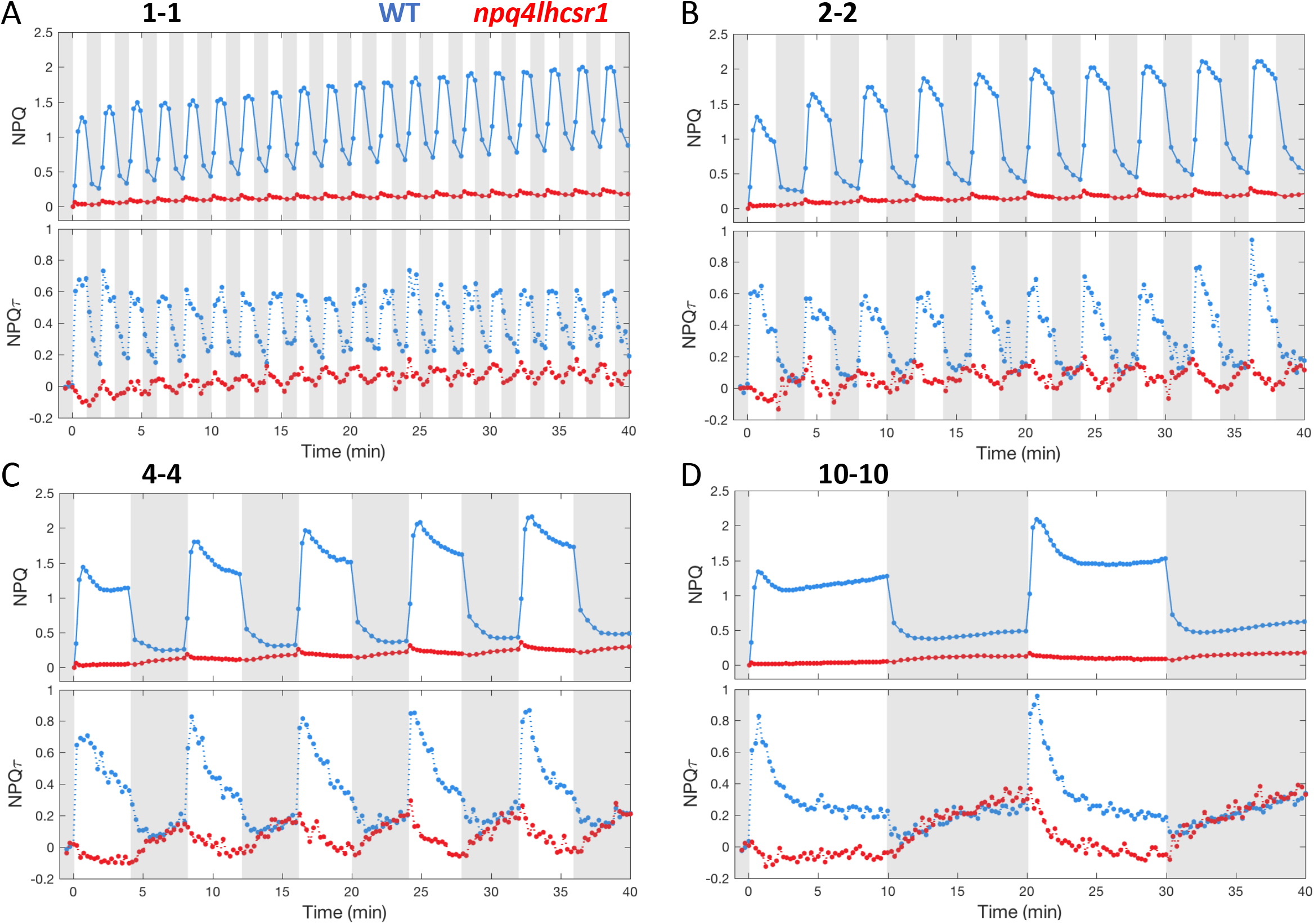
Quenching trajectories during light fluctuations in *npq4lhcsr1* and its control strain. The response of NPQ and NPQτ (upper and lower panel respectively) were measured in *npq4lhcsr1* mutant and its control strain (red and blue curves respectively) during 40 minutes of light fluctuations with periods of 1, 2, 4 and 10 minutes (**A, B, C** and **D** respectively) as described in **Fig. 1**. Shown are average of three biological replicates. For TCSPC data, each biological replicate was averaged from three technical replicates. The fluorescence lifetime values used to calculate NPQτ are shown in **Supp. Fig. 13**.

For the longer fluctuation periods ranging from 2 minutes to 10 minutes, after an initial burst of NPQ in HL, the level of NPQ decreased with continued illumination, eventually reaching a steady state for the 10 min fluctuating period (**Fig. 2**). A similar trend was also seen in NPQτ (**Fig. 2**). Therefore, these kinetics are directly related to chlorophyll fluorescence quenching, rather than other non-quenching processes that affect chlorophyll fluorescence. This phenomenon of decreasing NPQ during the light has been previously attributed to the consumption of the proton gradient by the activity of the CO_2_ concentration mechanism (CCM) (Burlacot *et al*., 2021). In addition to the effect arising from CCM, state transition via the induction of state 1 could also play some role in the NPQ decrease observed during HL periods, though this would not contribute during the first HL period in which cells are already in state 1 (following far-red acclimation). For fluctuating light periods longer than 4 minutes, upon a transition from HL to dark, the NPQ turned off rapidly but was then followed by a gradual rise in NPQ during further darkness, a trend that was also observed in NPQτ (**Fig. 2**). However, compared to NPQ, NPQτ showed a larger magnitude of increase during the long dark periods (**Fig. 2C, D**). Differences between the NPQ and NPQτ traces are considered in the discussion.

We conclude from these experiments that, when exposed to light fluctuations with periods ranging from 1 to 10 min, at least three components of NPQ are present: (i) a rapidly responding component, (ii) a slowly inducible component induced throughout the light fluctuations, and (iii) a component induced in the dark phases of light fluctuations.

### The majority of NPQ during light fluctuations is mediated by LHCSR proteins

It has been well established that LHCSR proteins are crucial for NPQ in *Chlamydomonas* during a single dark-to-light transition (Peers et al., 2009; Truong, 2011; Correa-Galvis *et al*., 2016). To examine the relative importance of each LHCSR protein for the functioning of NPQ during light fluctuations, we measured the chlorophyll fluorescence yield and lifetime during the same light-dark cycles on mutants impaired in the accumulation of LHCSR1 (*lhcsr1*) (Truong, 2011), LHCSR3-1 and LHCSR3-2 (*npq4*) (Peers et al., 2009), or all three LHCSRs (*npq4lhscr1*) (Truong, 2011; Ballottari et al., 2016). While the *npq4lhcsr1* mutant was highly impaired in its NPQ capacity for all light fluctuations (**Fig. 2**), single *npq4* and *lhcsr1* mutants showed some NPQ in response to light fluctuation (**Fig. 3**). Noticeably, for fluctuating periods longer than 4 minutes, the increasing NPQ observed during dark phases was more pronounced in the *npq4lhcsr1* mutant (**Fig. 2)**. We conclude from these experiments that although LHCSRs are responsible for most of the NPQ during the light phase of all light fluctuations, a substantial portion of the NPQ in WT is nonetheless mediated by other biochemical processes, part of which is induced during the dark periods of light fluctuations.

**Figure 3.**
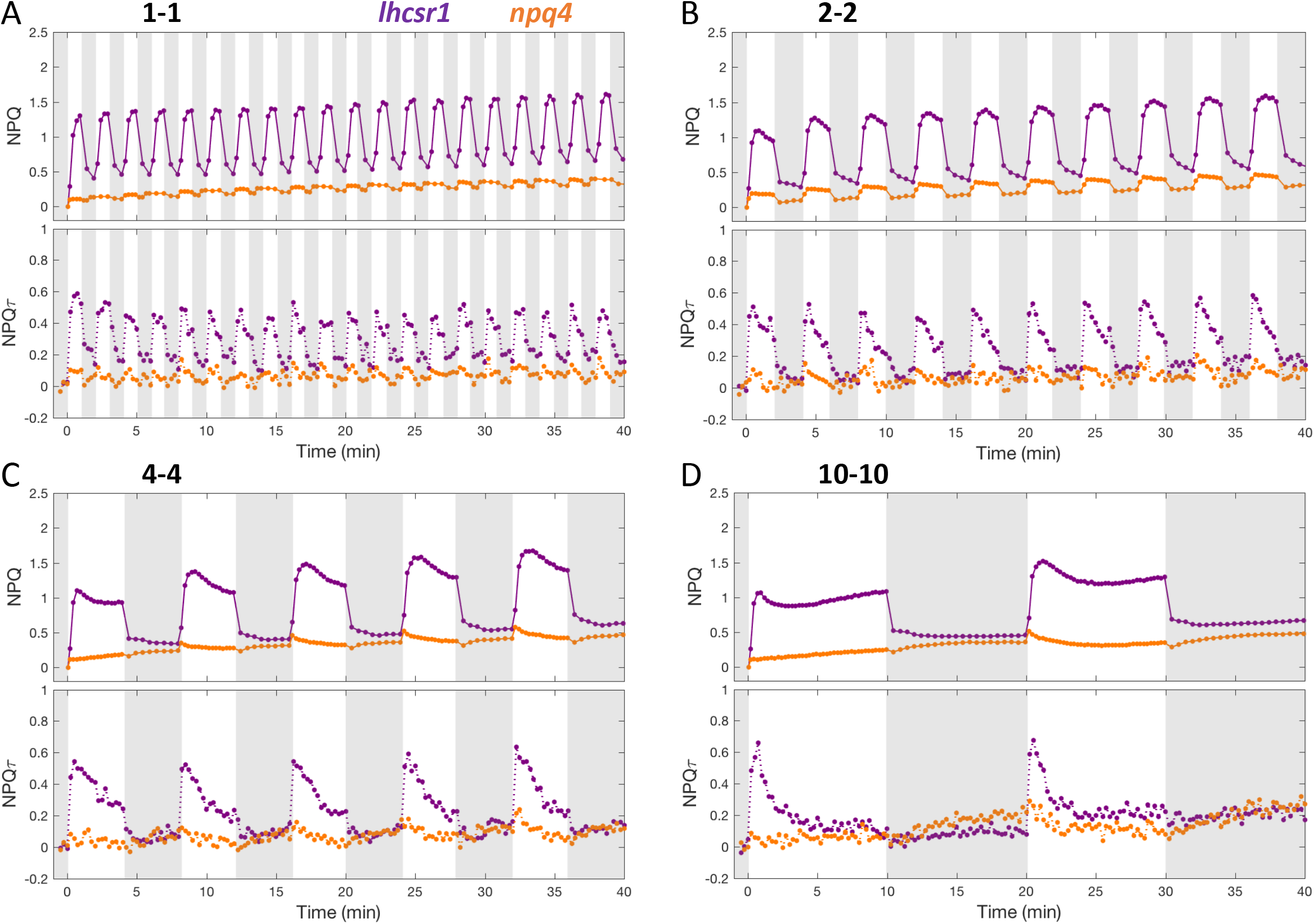
Quenching trajectories during light fluctuations in *lhcsr1* and *npq4*. The response of NPQ and NPQτ (upper and lower panel respectively) were measured in *lhcsr1* and *npq4* (purple and orange curves respectively) during 40 minutes of light fluctuations with periods of 1, 2, 4 and 10 minutes (**A, B, C** and **D** respectively) as described in **Fig. 1**. Shown are average of three biological replicates. For TCSPC data, each biological replicate was averaged from three technical replicates. The fluorescence lifetime values used to calculate NPQτ are shown in **Supp. Fig. 14**.

### The increasing quenching in the dark periods arises from state transition

Induction of NPQ during darkness has been previously reported in chlorophytes (Casper-Lindley and Björkman, 1996; Allorent et al., 2013), and qT has been proposed to be involved (Allorent et al., 2013). Re-organization of light-harvesting antennae between PSII and PSI was thus followed throughout a light fluctuation by measuring 77K fluorescence emission spectra. The spectra for cells were compared at three time points: after acclimation to far-red light (cells in state 1 (Zhang *et al*., 2021)), after 10 min HL, and after 10 additional min dark (see vertical lines in **Fig. 4A**). In WT and *npq4lhcsr1*, an increase in the emission at 710 nm specific to PSI-bound LHCII was observed between the 10 min (after HL) and 20 min (after dark) time points, suggesting that some re-association of LHCII from PSII to PSI occurs during the dark portion of our measurements (**Fig. 4, Supp Fig. 1**). In contrast, mutants lacking the STT7 kinase responsible for qT (*stt7* and *stt7npq4*) showed negligible changes in the 77K fluorescence emission spectra (**Supp Fig. 2**) and only showed a minimal increase in NPQ or NPQτ during the dark periods of light fluctuations (**Fig. 5**) and with increasing duration of exposure to light fluctuations (**Supp Fig. 3**).

**Figure 4.**
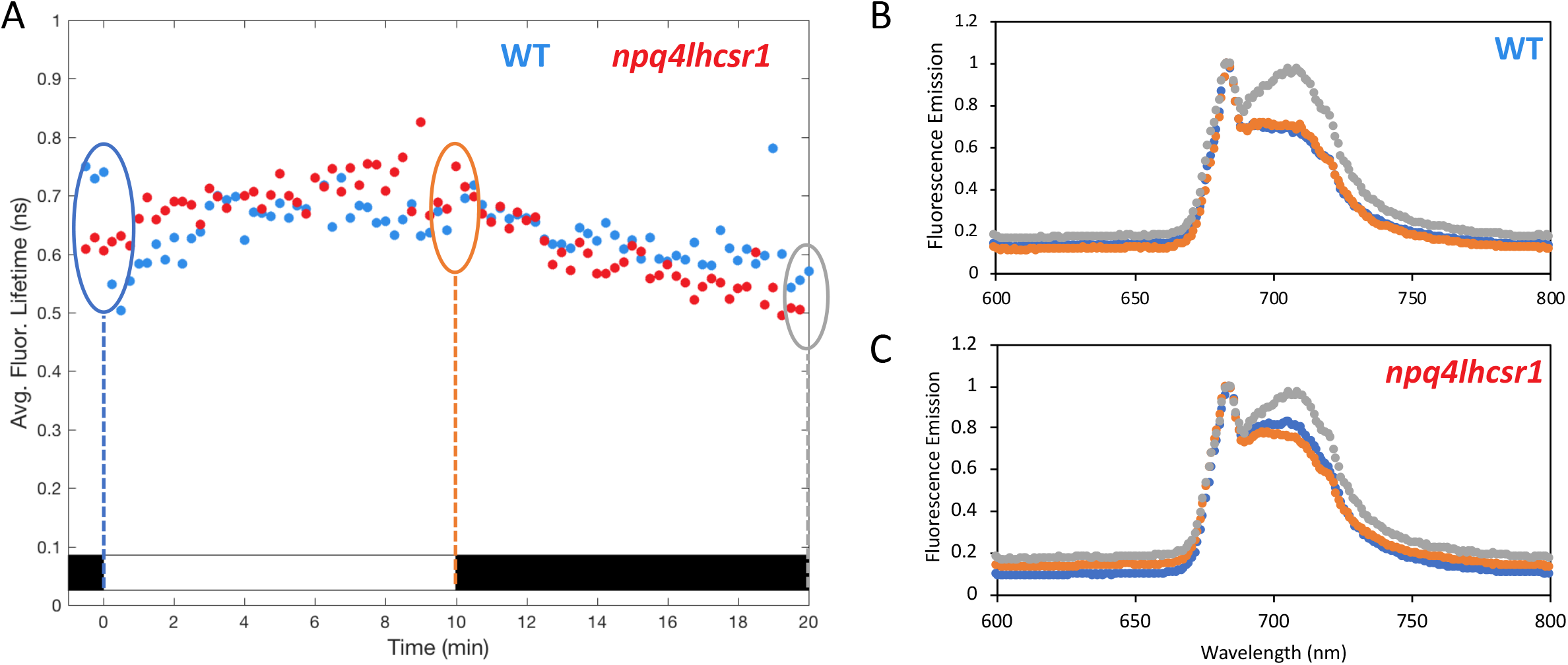
77K chlorophyll fluorescence emission spectra during the first high light-dark cycle of light fluctuations. Cells were placed in a TCSPC cuvette as described in **Fig. 1** and both fluorescence lifetime snapshots and 77K chlorophyll fluorescence emission spectra were taken through 10 minutes of high light and 10 minutes darkness. (**A**) Fluorescence lifetime trajectory of *npq4lhcsr1* mutant (red dots) and its control strain (WT, blue dots). On the graph, dashed vertical lines depict the timepoints at which samples were taken for 77K fluorescence spectra measurement. (**B, C**) 77K fluorescence emission spectra of samples taken in **A** on the control strain (WT, **B**) and *npq4lhcsr1* mutant (**C**). Spectra were taken at 0, 10, and 20 min timepoints (blue, orange and grey spectra respectively). Shown are representative spectra. Three independent biological replicate spectra for WT and npq4lhcsr1 are shown in **Supp. Fig. 1**. 77K spectra for the stt7 and stt7npq4 strains are shown in **Supp. Fig. 2**.

**Figure 5.**
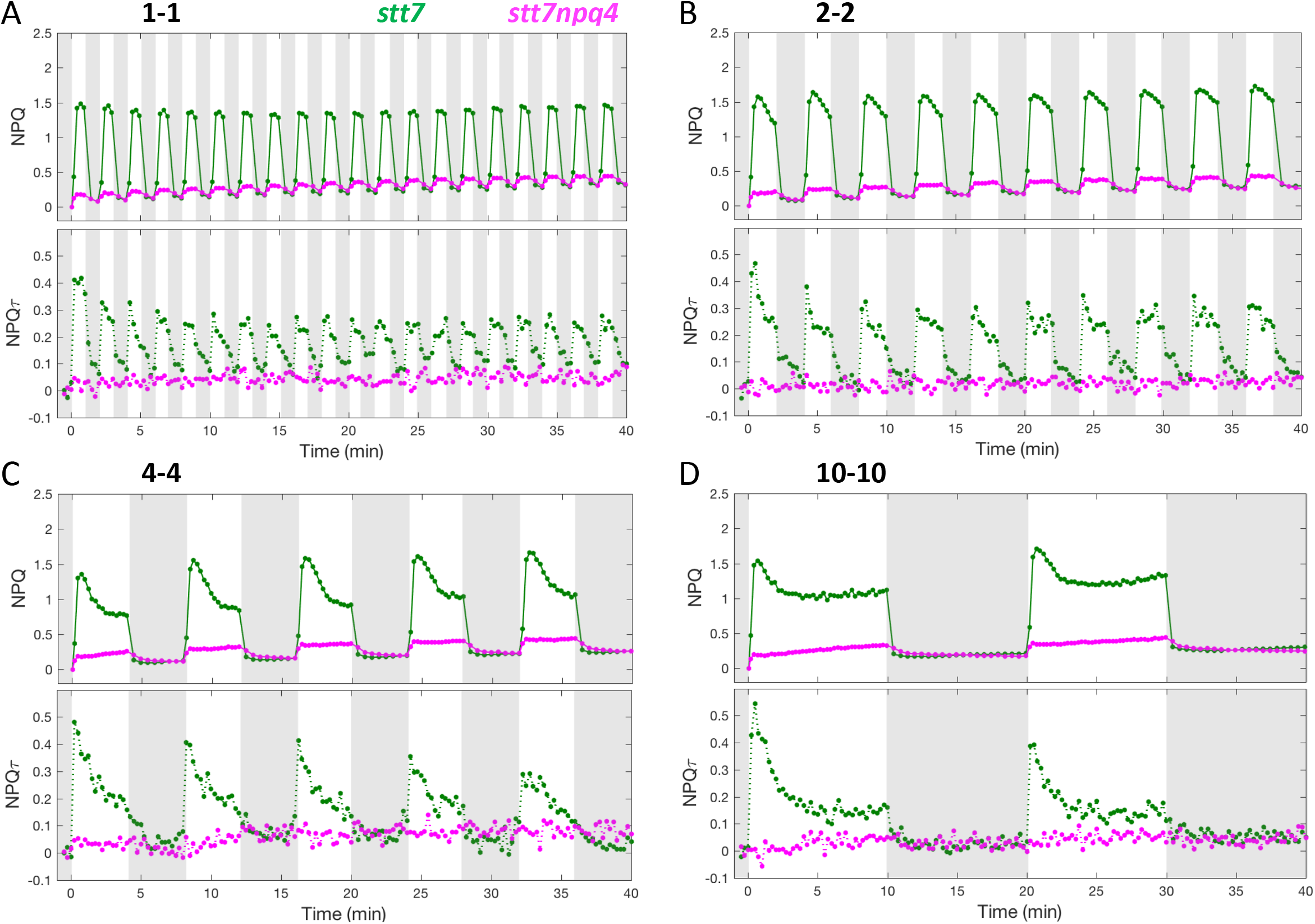
Quenching trajectories during light fluctuations in *stt7* and *stt7npq4*. The response of NPQ and NPQτ (upper and lower panel respectively) were measured in *stt7* and *stt7npq4* (green and magenta curves respectively) during 40 minutes of light fluctuations with periods of 1, 2, 4 and 10 minutes (**A, B, C** and **D** respectively) as described in **Fig. 1**. Shown are average of three biological replicates. For TCSPC data, each biological replicate was averaged from three technical replicates. The fluorescence lifetime values used to calculate NPQτ are shown in the **Supp. Fig. 15**.

To characterize the kinetics of qT occurring during the light-to-dark transition, we analyzed the response of the chlorophyll fluorescence lifetime for WT and *npq4lhcsr1* mutant cells that were exposed to 10 min of HL followed by 30 min of dark. Interestingly, upon light-to-dark transition, both strains showed steadily decreasing lifetimes for the first 10 min of darkness, after which, the fluorescence lifetimes began to reverse, eventually reaching the starting lifetime after 30 minutes of darkness (**Supp Fig. 4**). In contrast, *stt7* showed no such quenching of fluorescence lifetime during the 10 min dark period. Therefore, we conclude that the quenching observed during the dark phases of light fluctuations in *Chlamydomonas* arises from qT, which has an induction timescale of ∼10 min and is reversible upon continued exposure to darkness.

### Contributions of qE and qT to NPQ and growth under fluctuating light

It has been previously proposed that, while LHCSRs play an important role during short periods of illumination, state transitions are important for longer periods of high light acclimation (Erickson et al., 2015). To test this hypothesis and assess the scenario under fluctuating light conditions, we quantified the amount of NPQ that was mediated by each protein by comparing the remaining NPQ (or NPQτ) in each mutant relative to the NPQ (or NPQτ) in the WT reference strain (**Fig. 6A, Table 1**). Surprisingly, the contribution of each protein to overall NPQ did not seem to depend on the period of the light fluctuation (**Supp Fig. 5**). Therefore, as a first estimate for the contribution of LHCSRs and STT7, we calculated the average contribution under all four light fluctuation sequences. The donut charts in **Fig. 6 B, C, D** depict the contribution of each protein as a percentage of the total NPQ observed in the WT strain (represented as the complete donut). While LHCSR3 is responsible for the majority of overall NPQ (72%, **Fig. 6B**), STT7 had a substantial contribution mediating 42% of the NPQ, with LHCSR1 having a smaller contribution at 22% of NPQ. LHCSRs were found to have a substantially larger contribution during light phases, where they are responsible for 94% of WT NPQ, while in the dark phases, their contribution declined to 57% (**Fig. 6C, Supp Fig. 6**). On the other hand, STT7 played a significantly larger role in the NPQ during darkness (60%) than it does during illumination (36%) this latter result being likely due to inhibition of STT7 by high light (Rintamäki *et al*., 2000; Vink *et al*., 2004; Allorent et al., 2013). Interestingly, the amount of NPQ mediated by LHCSRs was more important during the beginning of light fluctuations while state transitions contributed more after 20 minutes of light fluctuation (**Fig. 6D, Supp Fig. 7**), revealing increased relative contribution of qT and decreasing contribution of qE with increasing time of exposure to light fluctuations.

**Figure 6.**
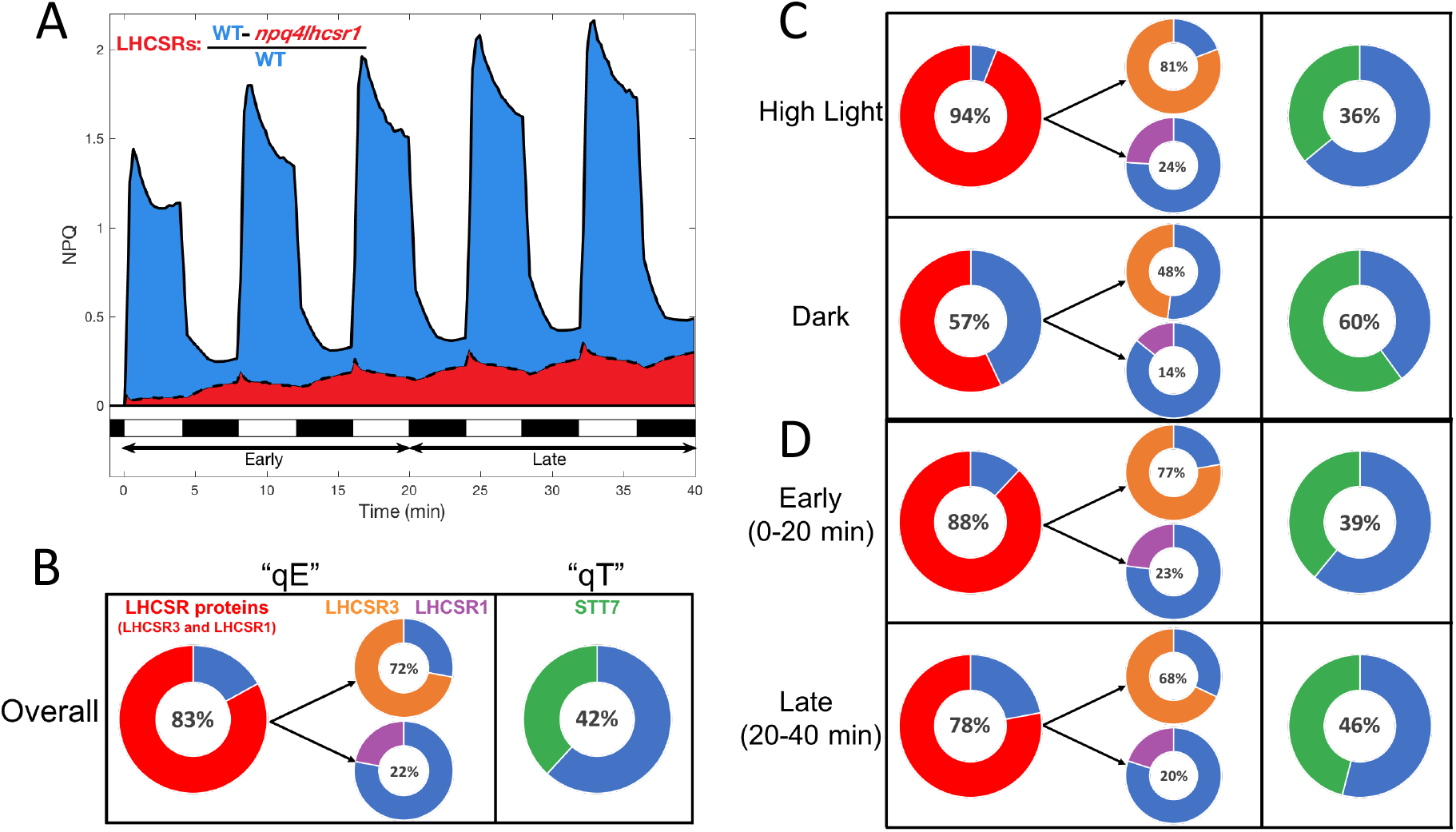
Quantification of the contribution of LHCSRs and STT7 to wild-type NPQ under fluctuating light. (**A**) Example of quantification of the relative NPQ mediated by LHCSRs. The area under the NPQ curve of *npq4lhcsr1* mutant (red) was subtracted from that of the control strain (blue) and expressed relative to the area of NPQ of the control strain. (**B, C**) Overall contribution of LHCSRs (red), LHCSR3 (orange), LHCSR1 (purple) and STT7 (green) averaged over all 40 minutes (**B**) or specific periods of the light fluctuations (**C**). For each protein, the part of the donut shown in red, orange, purple or green is the percentage of wild type NPQ lost in each mutant impaired in the accumulation of LHCSRs, LHCSR3, LHCSR1 and STT7 respectively. Given that the contribution of each protein was largely independent of HL/dark period (**Supp. Fig. 6**), shown are the average of all 4 light fluctuation sequences. Distribution of individual replicates and estimates of error (i) overall, (ii) during HL or dark and (iii) during early and late portion of light fluctuations are presented in **Supp. Fig. 5, 6, and 7, respectively**.

**Table 1.** Average contribution of each protein to overall wild-type NPQ for each light fluctuation sequence. Shown is the average value (n=6, evaluated from 3 TCSPC and 3 PAM replicates) and standard deviation of all individual replicates. The contributions of LHCSR3 (orange) and LHCSR1 (purple) were determined from the single mutants *npq4* and *lhcsr1*. The contribution of LHCSRs overall (red) was evaluated from the *npq4lhcsr1* mutant. The contribution of qT was assessed from the *stt7* mutant. Each error in the right column represents the standard deviation of each protein’s contribution across the 4 light fluctuation sequences. For simplicity, only the average values (shown in the right column) were used to generate **Figure 6** in the main text. Full distributions of the individual TCSPC and PAM data points are shown in **Supp. Fig. 5**.

| Overall Contributions | 1-1 | 2-2 | 4-4 | 10-10 | AVERAGE |
| --- | --- | --- | --- | --- | --- |
| LHCSR<sub>s</sub> | 89 ± 7% | 84 ± 7% | 83 ± 13% | 77 ± 24% | 83 ± 5% |
| LHCSR1 | 22 ± 12% | 22 ± 10% | 24 ± 16% | 19 ± 30% | 22 ± 2% |
| LHCSR3 | 81 ± 5% | 78 ± 5% | 73 ± 8% | 58 ± 15% | 72 ± 10% |
| STT7 | 46 ± 12% | 38 ± 16% | 46 ± 13% | 39 ± 22% | 42 ± 4% |

Since LHCSR3 is part of the mobile fraction of photosynthetic antenna and has been proposed to be affected by STT7 activity (Allorent et al., 2013), giving a potential mechanistic connection between qE and qT (Roach and Na, 2017), we aimed to interrogate the involvement of qT both together and separately from qE under fluctuating light. We compared the integrated PAM NPQ trajectories of *npq4stt7* relative to that of *npq4*, both of which lack LHCSR3, or *npq4stt7* relative to that of *stt7*, both of which lack STT7. The relative contribution of STT7 to NPQ was found to be different in the presence vs absence of LHCSR3, with STT7 playing a larger role when LHCSR3 is present (**Fig. 7, Table 2**). Likewise, the contribution of LHCSR3 was larger when STT7 was present. These data suggest that STT7 and LHCSR3 are cooperating to induce quenching during dark periods of light fluctuations. We propose that the activity of STT7 resulting in LHCSR phosphorylation and antenna mobility is responsible for this crosstalk between qE and qT during light fluctuations.

**Figure 7.**
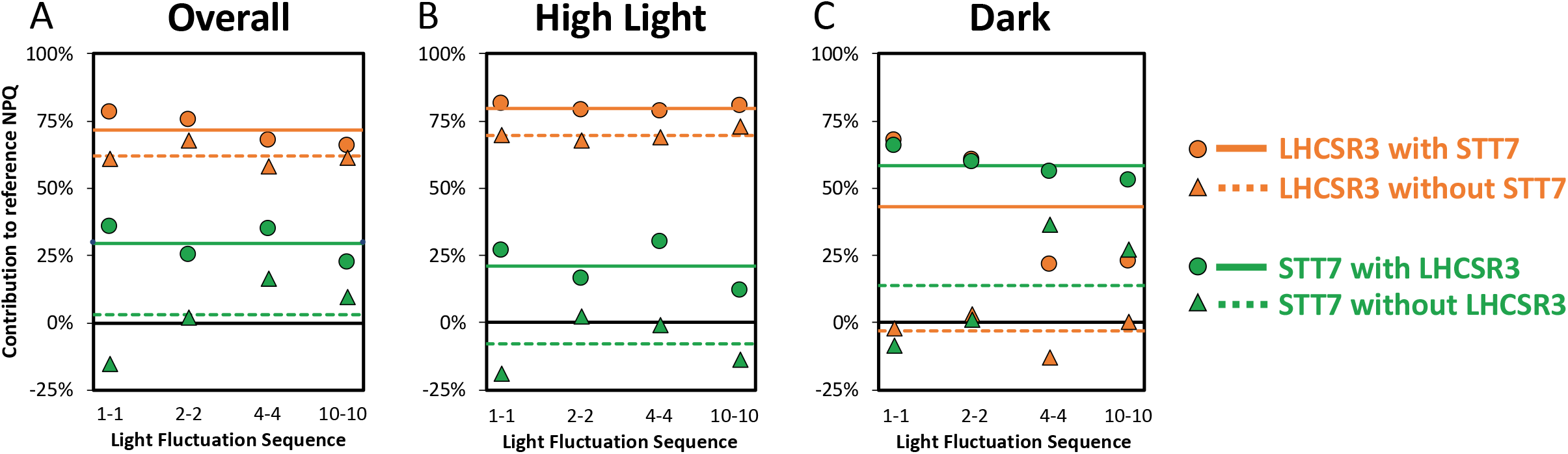
Untangling qE and qT. The quantification of the contributions of LHCSR3 (orange) and STT7 (green) to the NPQ response both in the presence (solid lines, circle markers) and absence (dashed lines, triangle markers) of the other protein is plotted as a function of the light fluctuation frequency. Horizonal lines, show the average contribution for each protein, considering all light fluctuation frequencies. For reference, the black lines represent zero contribution. The contribution of each protein was evaluated (**A**) over all 40 minutes of the experiment, (**B**) only during HL periods, or (**C**) only during dark periods. All of these were evaluated from PAM trajectories using various combinations of WT along with the *npq4, stt7*, and *stt7npq4* mutants as discussed in the Materials and Methods section.

**Table 2.** Average contribution of LHCSR3 and STT7 in the presence and absence of the other protein for each light fluctuation sequence. Shown is the average value (evaluated from 3 PAM replicates) and standard deviation of individual replicates. Each error in the right column represents the standard deviation of the 4 light fluctuation sequences. For simplicity, only the average values (shown in the right column) were used to generate **Figure 7A** in the main text.

| Overall Contributions | 1-1 | 2-2 | 4-4 | 10-10 | AVERAGE |
| --- | --- | --- | --- | --- | --- |
| LHCSR3 w/ STT7 | 78 ± 5% | 75 ± 5% | 67 ± 6% | 66 ± 14% | 72 ± 6% |
| LHCSR3 w/out STT7 | 61 ± 5% | 67 ± 4% | 58 ± 3% | 61 ± 3% | 62 ± 4% |
| STT7 w/ LHCSR3 | 36 ± 4% | 25 ± 11% | 35 ± 8% | 22 ± 9% | 29 ± 7% |
| STT7 w/o LHCSR3 | -16 ± 15% | 2 ± 12% | 16 ± 5% | 9 ± 6% | 2 ± 14% |

Although mutants impaired in the accumulation of LHCSRs and STT7 have strongly impaired NPQ capacities, this does not seem to impair the growth of those strains under continuous high light conditions (Depege et al., 2003; Peers et al., 2009; Cantrell and Peers, 2017). Recent data have shown that the growth of *npq4* and *npq4lhcsr1* mutants is impaired under certain light fluctuation conditions (Cantrell and Peers, 2017; Roach, 2020). We therefore investigated whether the growth impairment of those strains could be dependent on the period of light fluctuation. While all the mutants grew as well as the control strain under continuous illumination (**Supp. Fig. 8**), *npq4, lhcsr1, npq4lhcsr1* and *stt7npq4* mutants exhibited impaired growth under fast light fluctuations with a 1-minute period (**Fig. 8**). In contrast, only *npq4lhcsr1* and *stt7npq4* mutants showed an impaired growth under slower light fluctuations with a period of 10 minutes, and the growth of all mutants was similar when the period was increased to 30 minutes (**Fig. 8**). We conclude from this experiment that qE mediated by LHCSR proteins is critical for growth under light fluctuations and that this role is more important for rapid light fluctuations.

**Figure 8.**
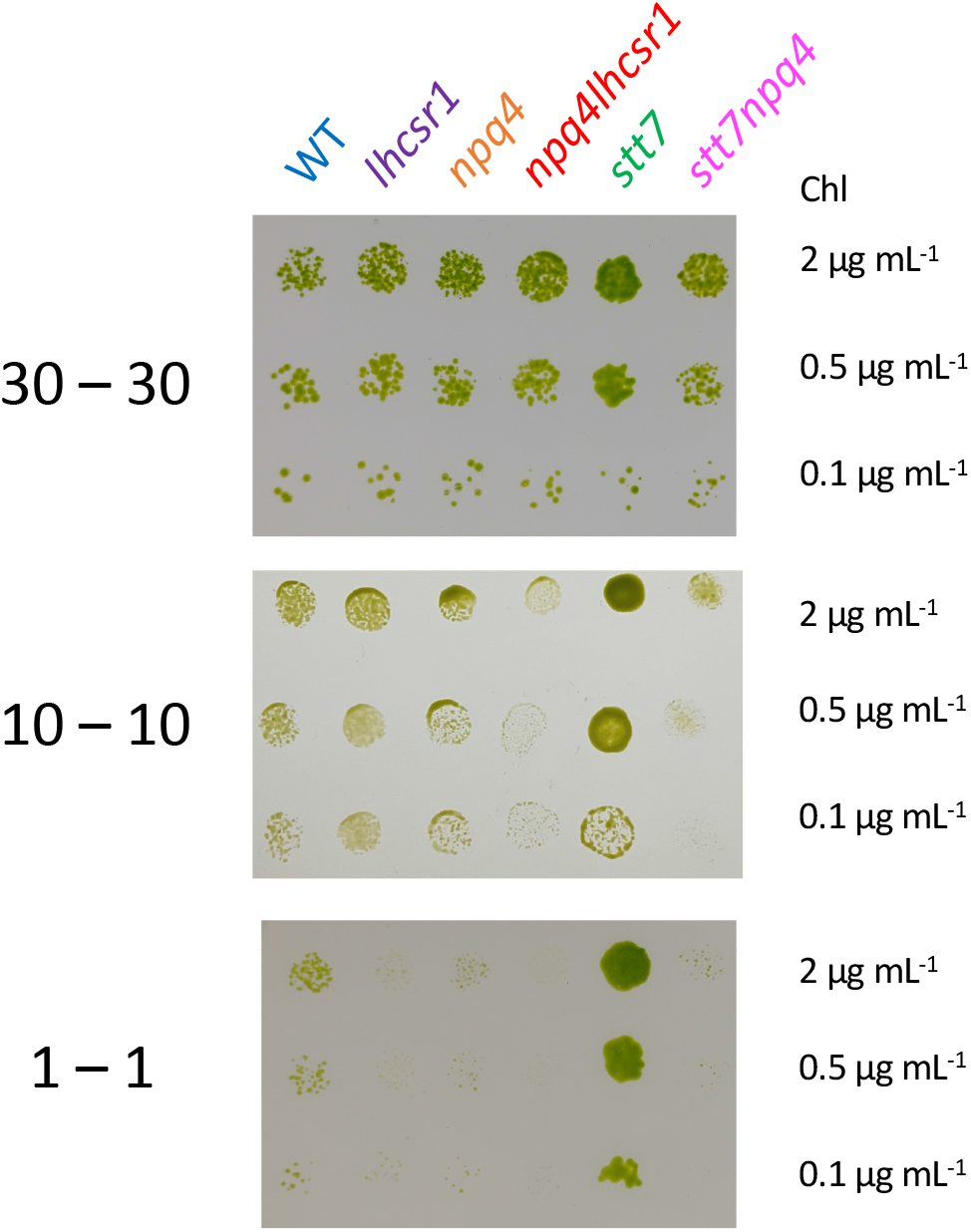
Growth of mutants impaired in qE and/or qT under various periods of dark/light cycles. *lhcsr1, npq4, npq4lhcsr1, stt7* and *stt7npq4* mutants and their control strain (WT) were diluted and spotted at different chlorophyll concentration and grown on plates under dark/light cycles with a period of 30 (30-30, upper panel), 10 (10-10, middle panel) or 1 minute (1-1, lower panel). Each row represents a different chlorophyll concentration. Shown are representative spots of three biological replicates. Growth under constant low light or high light are shown in **Supp. Fig. 8**.

## Discussion

The involvement of pH-sensing LHCSR proteins and state transitions in the photoprotective response of *Chlamydomonas* has been previously described (Peers et al., 2009; Allorent et al., 2013). While mutants impaired in accumulation of LHCSRs were shown to have limited growth when light is provided in a day/night cycle (Cantrell and Peers, 2017) or fluctuating with a 10-minute period (Roach, 2020), our understanding of the contribution of LHCSRs and state transitions to photoprotection during fluctuating light remains limited. Here, by measuring the NPQ levels during light fluctuations in a range of mutants impaired in the accumulation of LHCSR3, LHCSR1, and/or STT7 we have unraveled their relative contributions to NPQ. Varying the length of fluctuating periods from 1 to 10 min allowed us to assess the dynamics of rapid qE- and slower qT-type processes. Interestingly, we observe that qT builds up during the dark periods of the light fluctuations and continues to play a role in the NPQ response during subsequent light phases. It occurs even in the absence of LHCSR3 (see *npq4* and *npq4lhcsr1* mutant in **Fig. 2** and **Fig. 3**), has a timescale of 10 min, and is reversible (**Supp Fig. 4**), which is consistent with recent literature (Allorent et al., 2013; Zhang et al., 2021). During a transition between low-light and high-light stress, qE proteins take a few hours to be fully induced (Peers et al., 2009), and it was hypothesized that qT may substitute for qE during this delay (Allorent et al., 2013). Our results show that even when LHCSRs are fully activated (i.e., in high light-acclimated cells), the occurrence of qT remains substantial during light fluctuations (**Fig. 2**). State transition or qE mutants were previously shown to have high reactive oxygen species (ROS) production (Allorent et al., 2013; Barera *et al*., 2021). Thus, the substantial amount of qT induced during the dark periods of light fluctuations may enhance photoprotection and limit ROS production by “anticipating” the next exposure to high light. The combination of fast qE (turns on rapidly upon HL exposure due to ΔpH) and residual qT (from previous dark periods) could therefore provide effective photoprotection in an unpredictable fluctuating-light environment.

Since qE has long been ascribed as the fastest component of NPQ, directly responding to the thylakoid lumen proton concentration (Briantais et al., 1979), and qT as a slower component (Allorent et al., 2013), the contribution of qE to NPQ was proposed to be stronger for short periods of HL, with the contribution of qT becoming increasingly important during longer periods of high-light stress (Erickson et al., 2015). Our approach of systematically assessing the response of photosynthesis to various periods of light fluctuations has revealed nuances in this interpretation. Surprisingly, we found that the overall contributions of qE and qT are different in the presence vs absence of the other component (**Fig. 7**) revealing that interactions between qE and qT contribute to the NPQ response during all periods of light fluctuation. Additionally, the *npq4* mutant showed significantly reduced NPQ capacity compared to WT in the dark periods (by 74%) under fast light fluctuations (1-1 sequence), but only a 17% impairment under slower fluctuations (10-10) (see **Supp. Fig. 6b**), showing that relaxation of qE (around 1 min) is mediated by LHCSR3 and contributes substantially to the response of NPQ to short periods of light fluctuations. Such relaxation kinetics may contribute to a faster response of NPQ to the next illumination if the period of light fluctuations is shorter than 2 minutes. It is also worth noting that the relative importance of qE in NPQ decreased after 20 minutes of light fluctuations, while the opposite occurred for qT (**Fig. 6**), reflecting a build-up in qT throughout the 40 minutes of light fluctuations. Modeling the response of photosynthesis to complex light fluctuations has been done (Zaks *et al*., 2012; Zaks *et al*., 2013; Tanaka *et al*., 2019; Steen *et al*.; Nedbal and Lazár, 2021) and would allow targeting specific mechanisms for increasing plant yields in the field. Our results show that such efforts should consider both the period of light and dark as well as the total time exposed to fluctuating light. The light intensities used during light fluctuations may also affect the relative contributions of each mechanism. In green microalgae it is tempting to speculate that in nature, where exposure to HL and dark occur repeatedly, qE may play a more important role in the beginning of light fluctuations while qT may provide a photoprotective response on a longer time scale.

Although qI is known to contribute to NPQ during prolonged illumination, it is likely negligible in our experimental conditions since both *npq4lhcsr1* and *npq4stt7* mutants did not show a strong buildup of NPQ or NPQτ during the 20 min of HL exposure during the light fluctuations (**Fig. 2** and **Fig. 5**). Additionally, Fv/Fm were not significantly different between strains prior to light fluctuations, with the exception of *npq4lhcsr1* which showed a lower Fv/Fm, reflecting some constitutive qI following overnight HL acclimation in the complete absence of LHCSRs (**Supp Fig. 9**).

Interestingly, when comparing NPQ and NPQτ, the magnitude of the quenching decrease during HL was larger for NPQτ than NPQ (**Fig. 2**). The energetic requirement (and thus its proton gradient consumption) of the CCM depends on the inorganic carbon (C_i_) availability (Fei *et al*., 2021). Since the high cell concentration in the TCSPC sample leads to strong C_i_ consumption, this could deplete the C_i_ concentration even in the presence of bubbling, leading to a C_i_ level sensed by cells in the TCSPC sample being lower than what is experienced by cells in the PAM sample. This would lead to higher activity of CCM, and hence a larger decrease in quenching, under TCSPC sample conditions compared to PAM sample conditions. The decrease in both NPQ and NPQτ was also stronger for longer HL periods as well as later in the sequence (**Fig. 2**). The slope of the initial decrease in NPQτ or NPQ during HL was similar for all four sequences (**Supp. Fig. 10**) and is likely dictated by C_i_ availability and its influence on the CCM kinetics. The magnitude of the decrease in NPQ and NPQτ was larger for longer light periods (**Supp. Fig. 10d**,**h**), likely due to simultaneous activation of the CCM that dissipates the proton gradient and the onset of slower forms of NPQ such as state transitions. Conversely, the differences in the magnitude of the increase in NPQ and NPQτ during the dark periods could be due to differences in O_2_ concentrations sensed by the cells in both conditions, which is known to affect the extent and rate of state transition (Forti and Caldiroli, 2005).

Although LHCSR3 plays the dominant role in photoprotection under constant and fluctuating light conditions, we also observe a role for LHCSR1 in our measurements. While the chlorophyll fluorescence dynamics of *lhcsr1* are similar to those of WT **(Fig. 3)**, we observe a ∼20% reduction in overall NPQ in this mutant under fluctuating light conditions **(Fig. 6)**. This small amount of photoprotection afforded by LHCSR1 *in vivo* is consistent with previous *in vitro* investigations (Dinc *et al*., 2016; Nawrocki *et al*., 2020) in which LHCSR1 has been suggested to mediate energy transfer between LHCII and PSI (Kosuge *et al*., 2018) and to compensate for the absence of LHCSR3 (Girolomoni et al., 2019). Interestingly, our results suggest that different from the case for LHCSR3, the qE that is mediated by LHCSR1 is largely frequency independent (**Supp. Fig. 5**). Both LHCSR1 and LHCSR3 are thought to generate NPQ in response to (i) the proton gradient and (ii) carotenoid composition (Kondo et al., 2017; Troiano et al., 2021), thus the frequency-dependent LHCSR3 and frequency-independent LHCSR1 could differ in their pH or carotenoid dependency.

The relationship between NPQ capacities and growth have remained puzzling in *Chlamydomonas*, because only some light fluctuation regimes have consistently been shown to impair growth (Peers et al., 2009; Truong, 2011; Cantrell and Peers, 2017; Roach, 2020). Here we show that all mutants lacking LHCSRs showed impaired growth under rapid light fluctuations (1-1 sequence), and that this impairment was lower under slower fluctuations (10-10 sequence) and absent under an even slower 30-30 sequence or constant illumination (**Fig. 8** and **Supp. Fig. 8**). There seems to be a good relationship between defect of NPQ and growth deficiency under short time-scale fluctuations when considering *npq4lhcsr1* and *stt7npq4* mutants (**Fig. 8**), clearly showing that LHCSR-dependent qE is critical for growth when light fluctuates with short period of time. However, surprisingly, the growth defect of *lhcsr1* seemed larger than *npq4* under the 1-1 sequence. This suggests that the growth capacity of LHCSR mutants under light fluctuations may not depend only on the level of NPQ. The growth defects observed for qE mutants under short time-scale light fluctuations may be affected by deactivation of the Calvin-Benson-Basham (CBB) cycle during the dark phases, as previously proposed by (Roach, 2020). Such a frequency dependence for the growth may thus additionally reveal the specific time scales of CBB cycle activation and deactivation during extended periods of fluctuating light. Other factors may include activation of compensatory mechanisms that enable photoprotection at the expense of growth or the increased production of reactive oxygen species in *npq4* mutants (Roach *et al*., 2020; Barera et al., 2021), which could be greater in the *lhscr1* mutant. Note here that in our conditions, due to slightly different genetic background between *stt7* and the WT control, we cannot make a conclusion on the mechanism by which *stt7* grew better under short timescale fluctuations (**Fig. 8**). The WT background may be particularly sensitive to fast 1-1 and 10-10 fluctuations; another possibility could be that qT is detrimental for growth under medium to short time scale fluctuations.

Through 77K fluorescence emission spectra analysis, we have shown that the increasing dissipation observed in the dark requires the STT7 kinase responsible for state transition **(Fig. 4** and **Supp. Fig. 2)**. This effect, already described in *Chlamydomonas* (Allorent et al., 2013) and *Dunaliella salina* (Casper-Lindley and Björkman, 1996), greatly contributes to NPQ during light fluctuations. In plants, the occurrence of state transitions and its involvement in NPQ is thought to be minor (Allen, 1992; Minagawa, 2011) even if mutants of *Arabidopsis thaliana* impaired in state transition (*stn7*) exhibit impaired growth under light fluctuations (Bellafiore *et al*., 2005). Interestingly, an increase of NPQ during the dark periods of fluctuating light was recently reported in *npq4* leaves of *A. thaliana* (Steen et al., 2020) for which about 53% of the WT NPQτ remained in the mutant after 40 minutes of exposure to light fluctuations despite the absence of the pH-sensing protein PsbS (**Supp. Fig. 11**). However, it remains unclear as to how much of the dark quenching in plants originates from qT as opposed to effects related to de-epoxidized xanthophyll pigments (Steen et al., 2020) or LHC protein conformation and/or aggregation (Goral *et al*., 2012). In the future, periodic illumination experiments performed on *A. thaliana* mutants impaired in qE and/or qT could clarify the relative importance of each mechanism and allow a comparison of the *in vivo* functioning of NPQ in higher plants and green algae under light fluctuations.

Tuning the relaxation kinetics of NPQ in higher plants has been shown to improve crop plant productivity (Kromdijk *et al*., 2016), and recent modelling of the response of plant canopies to natural light fluctuations has shown that there remains ample room for improving photosynthetic efficiency under non-steady state conditions (Wang *et al*., 2020). Similar opportunities for improvement also exist for increasing biofuel production from microalgae (Benedetti *et al*., 2018; Perin and Jones, 2019; Vecchi *et al*., 2020). In both cases, such optimization will require detailed understanding of the dynamic activity of NPQ mechanisms. The sum of the NPQ contributions of each protein is greater than 100% (**Fig. 6**) which cannot be explained by a difference in the amount of LHCSR3 between WT and *stt7* (**Supp. Fig. 12**). Additionally, our quantifications of qE and qT reveal that each of them depends on the presence of the other process (**Fig. 7**), showing that there is an interaction between qE and qT in *Chlamydomonas* under fluctuating light. The possibility of an interaction between qE and qT has been previously suggested on the basis of a kinetic analysis of qT in the presence and absence of LHCSR3 protein (*npq4* mutant) (Roach and Na, 2017). Our findings are consistent with a partial overlap of the functions of LHCSR3 and STT7 in both qE and qT. This partial overlap highlights the need for further investigations of the interactions occurring between proteins that underlie the *in vivo* NPQ response, not only in HL but also in dark, and more generally during light fluctuations. For example, in microalgae, the quenching mediated by LHCSR3 both at PSII and PSI level (Girolomoni et al., 2019) could be tuned by the movement of LHCSR3 from PSI to PSII during state transitions (Allorent et al., 2013). Thus, the described LHCSR3 phosphorylation by STT7 (Bergner *et al*., 2015) could explain the crosstalk between qE and qT. LHCSR3 is also known to associate with LHCII trimers in the PSII supercomplex (Semchonok *et al*., 2017); therefore, a similar LHCSR3-LHCII interaction may also generate quenching in the trimers following the detachment of LHCII from PSII. It should be noted that in the thylakoid membrane, LHCII can exist in a range of different conformations and/or quenching states (Tian *et al*., 2015; Kawakami *et al*., 2019). At this point it is not possible to distinguish the relative contributions of different forms of LHCII in individual snapshot measurements. The ensemble fluorescence lifetime likely originates from some combination of at least three LHCII subpopulations: unphosphorylated and bound to PSII (state 1) (Drop *et al*., 2014) with an intermediate fluorescence decay component, phosphorylated and unbound (free LHCII) (Iwai *et al*., 2010b) which has been previously assigned to a long fluorescence decay component (Ünlü et al., 2014), and phosphorylated and bound to PSI (state 2) (Huang *et al*., 2021) with a short fluorescence decay component. In intact algal cells, the relative abundance of each form of LHCII likely dynamically evolves throughout exposure to the fluctuating HL-dark sequences. Further development of *in vivo* spectroscopic tools will be required to disentangle the dynamics of LHCII conformations and correlate them with photoprotection. Overall, a deeper understanding of the protein interactions underlying NPQ dynamics will be highly valuable in finding new ways to improve plant and microalgal productivity.

## Conclusions

LHCSR- and STT7-mediated nonphotochemical quenching processes (qE and qT) are known to underly the photoprotective response of the microalgae *Chlamydomonas*. Here, we have applied a new method to disentangle the involvement of qE and qT in real time by exposing intact algal cells to repetitive cycles of high light and darkness alternating at different frequencies. While both qE- and qT-type responses are present during all light fluctuations, LHCSR-dependent qE plays a larger role in the beginning of light fluctuations and during the HL periods. The contribution of STT7-dependent qT became more pronounced upon longer exposure to fluctuating light and especially during the dark periods of light fluctuations. Over the long term, rapid light fluctuations reduced the growth of mutants impaired in LHCSRs, demonstrating the importance of LHCSR proteins during abrupt changes in light intensity. Overall, our work shows that a cooperativity between LHCSR proteins and STT7 constitutes an important regulatory feature of the photoprotective response in *Chlamydomonas*, likely mediated by phosphorylation of LHCSR3 by STT7. These findings provide a foundation for disentangling and modelling how the diverse molecular mechanisms involved in plant and microalgal acclimation to light fluctuations interact and enable robust photosynthesis in nature. We envision that further knowledge on the response of photosynthetic mechanisms to various periods of light fluctuations will open new avenues for building a strong understanding of how photosynthetic organisms respond to complex light fluctuations.

## Accession numbers

Genes studied in this article can be found on https://phytozome.jgi.doe.gov/ under the loci Cre08.g365900.t1.2 (LHCSR1), Cre08.g367500.t1.1 (LHCSR3.1), Cre08.g367400.t1.1 (LHCSR3.2), Cre02.g120250.t1.1 (STT7).

## Supporting information

Supplemental informations

## List of abbreviations

NPQ: non-photochemical quenching
qE: energy-dependent quenching
qT: state transitions
qI: photoinhibition
qZ: zeaxanthin-dependent quenching
PAM: pulse-amplitude modulation
TCSPC: time-correlated single photon counting
LHCSR: light-harvesting stress related protein
STT7: serine/threonine-protein kinase
LHC: light-harvesting complex
PS: photosystem
CCM: CO_2_ concentration mechanism
C_i_: inorganic carbon (CO_2_, HCO_3_^-^, CO_3_^2-^)
HL: high light

## Materials and Methods

### Strains and culture conditions

*Chlamydomonas* mutants and their respective wild-type 4A-were previously described (*npq4* (Peers et al., 2009), *lhcsr1* (Truong, 2011), *npq4lhcsr1* (Truong, 2011), *stt7-9* (Cardol et al., 2009), *stt7npq4* (Allorent et al., 2013)). All strains were grown photoautotrophically under moderate light (50 µmol photons m^-2^ s^-1^) in minimal HS medium under air level of CO_2_ (20°C). Except for the growth test, cell cultures (5-8 µg Chl mL^-1^) were incubated overnight at high light (400 µmol photons m^-2^ s^-1^), for maximizing expression of LHCSR proteins (Tibiletti *et al*., 2016). Prior to each measurement, cells were illuminated for at least 15 min with low intensity far-red light (3in1LED panel with far-red LED; 3LH series, NK system, Japan) to ensure a complete state 1 configuration (Bonaventura and Myers, 1969). All replicates shown are biological replicates from independent cultures.

### Chlorophyll Fluorescence Measurements

In this work, we employ two techniques to monitor the activation and deactivation of NPQ throughout 40 minutes of exposure to repeated periods of high light and dark on the basis of changes in Chl fluorescence emission (see **Fig. 1** for an illustration of the experimental design). Time-resolved Chl fluorescence was measured via time-correlated single photon counting (TCSPC) while Chl fluorescence yield was measured in parallel experiments using pulse-amplitude modulation (PAM) fluorimetry. Although both methods can monitor NPQ, the fluorescence lifetime is not susceptible to a range of non-quenching processes that can impact the fluorescence yield (such as chromophore bleaching, changes in chlorophyll concentration, chloroplast movement, or enhanced light scattering (Zaks et al., 2013; Sylak-Glassman *et al*., 2016). Therefore, fluorescence lifetime measurements provide insight into processes that directly quench chlorophyll fluorescence.

#### A) TCSPC measurement and fitting

The average chlorophyll fluorescence lifetime was measured by time-correlated single photon counting (TCSPC), as previously described (Sylak-Glassman et al., 2016; Steen et al., 2020). A diode laser (Coherent Verdi G10, 532 nm) pumped a Ti:Sapphire oscillator (Coherent Mira 900f, 808 nm, 76 MHz) and the output was subsequently frequency doubled using a β-barium borate crystal to obtain 404 nm light. These pulses were used for excitation of the sample with a power of 1.7 mW (20 pJ/pulse) and Chl fluorescence emission at 680 nm was detected via an MCP-PMT (Hamamatsu R3809U). A custom-built LABVIEW software was used to synchronize three shutters located in the laser path, actinic light path, and the path between the sample and detector. Every 15 sec, a fluorescence lifetime snapshot measurement was acquired by exposing the cells to the saturating laser (404 nm) for 1 second and detecting the emission. Fluorescence lifetime snapshots were measured by TCSPC using a Becker-Hickl SPC-850 data acquisition card and SPCM software. In between the snapshot measurements, high-light illumination of the cells was achieved by exposing the cuvette to white light set to an intensity of 620 μmol photons m^-2^ s^-1^ (Leica KL1500 LCD, peak 648 nm, FWHM 220 nm), chosen as the minimal light level necessary to saturate photosynthetic reactions and induce qE activity at the onset of illumination. It was also chosen such that PAM and TCSPC would use identical high light intensities. The sample concentration was adjusted to ∼80 µg Chl mL^-1^ for TCSPC measurements. To control the gas composition of the culture and prevent cells from settling to the bottom of the cuvette, the sample was bubbled with air (ambient CO_2_ concentrations) at a rate of 2-7 mL min^-1^ throughout the entire 40 min experiment duration, although note that such bubbling increased the noise of the measurements.

For each fluorescence decay measurement, to ensure that PSII reaction centers were closed, we selected the 0.2 s step with the longest lifetime from the overall 1 s snapshot measurement duration (Sylak-Glassman et al., 2016). This longest step was then fit to a bi-exponential decay (Picoquant, Fluofit Pro-4.6) and the average amplitude-weighted fluorescence lifetime (τ_avg_) was calculated for each snapshot measurement. The NPQτ parameter is derived from the fluorescence lifetime snapshot measurements and is defined analogously to NPQ (Sylak-Glassman et al., 2014; Sylak-Glassman et al., 2016): 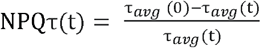. The value of NPQτ represents the magnitude of the quenching response based on the change in the average fluorescence lifetimes between time t=0 (after far-red acclimation but before HL exposure) and any other time *t* during the 40 min snapshot trajectory. Therefore, using NPQτ removes confounding effects arising from any differences in the average chlorophyll excited state lifetime of the different strains following far-red acclimation. For all TCSPC measurements, each biological replicate represents the average of three technical replicates measured on the same day.

#### B) PAM measurements

Chlorophyll fluorescence yield was measured using a pulsed-amplitude modulation (PAM) fluorimeter (Dual-PAM 100, Walz GmbH, Effeltrich, Germany) with the red measuring head. Red saturating flashes (8,000 µmol photons m^−2^ s^−1^, 600 ms, 620 nm) were delivered to measure F_M_ (maximal fluorescence yield in dark-acclimated samples) and then every 15 s or 30 s to measure F_M_′ under actinic light exposure or dark phase respectively. Actinic light illumination (620 nm) was set to 620 μmol photons m^-2^ s^-1^. Fluorescence emission was detected using a long-pass filter (>700 nm). NPQ was calculated as (F_M_ − F_M_′)/F_M_′. The Chl concentration was ∼5-8 µg Chl mL^-1^ and as for TCSPC, all PAM measurements and the sample was bubbled with air at a flux of 2-7 mL min^-1^ for proper control of the gas concentrations of the sample throughout the entire 40 min experiment duration, note that such bubbling increased the noise of the measurements (but to a lesser extent than for TCSPC).

#### C) Quantifying the contributions of LHCSRs and STT7 to NPQ

To assess the relative contributions of LHCSR1, LHCSR3, and STT7 to overall NPQ, we analyzed the NPQ (PAM) and NPQτ (TCSPC) trajectories for WT and each mutant. The relative contribution of each protein was determined as the percent change in the integrated snapshot trajectory of NPQ or NPQτ for each mutant relative to the control WT strain. As the contribution of each actor was found to be overall independent of HL-dark fluctuation frequencies in the range of 1 min^-1^ to 0.1 min^-1^. (**Supp. Fig. 5**), the average contribution of each protein under the four light fluctuating sequences for both PAM and TCSPC was considered. Additionally, to characterize the involvement in activation or deactivation of NPQ, the quenching trajectories were integrated solely under HL or dark periods, respectively (**Supp. Fig. 6**). The contributions of each protein to the early vs. late responses were further assessed by integrating from 0-20 min and 20-40 min, respectively (**Supp. Fig. 7**). These results are summarized in **Fig. 6** of the main text. Since it has been suggested that the *stt7-9* mutant may be “leaky” (Bergner et al., 2015), our estimates for the STT7 contribution are lower-bound estimates. The potential leakiness of the *stt7-9* mutant does not affect any of the conclusions presented in the manuscript. In addition to quantifying the relative contribution to NPQ of each protein relative to the WT strain, we also assessed the contribution of STT7 in the absence of LHCSR3 by integrating the PAM NPQ trajectory of *npq4stt7* relative to that of *npq4*. Likewise, the contribution of LHCSR3 was assessed independently of STT7 by integrating the PAM NPQ trajectory of *npq4stt7* relative to that of *stt7*.

### 77K Chlorophyll fluorescence emission

Chlorophyll fluorescence emission spectra of *Chlamydomonas* cells at 77 K were obtained by freezing whole cells (∼5-8 µg Chl mL^-1^ final concentration) in liquid nitrogen. The emission spectrum was then measured between 600 and 800 nm (435 nm excitation wavelength, RF-5300PC spectrophotometer, Shimadzu).

### Growth tests

The different *Chlamydomonas* strains were cultivated photoautotrophically under moderate light (50 µmol photons m^-2^ s^-1^) in minimal medium under air level of CO_2_ (20°C). Cells were harvested during exponential growth and resuspended in fresh minimal medium to 0.1, 0.5, or 2 µg Chl mL^-1^. Eight-microliter drops were spotted on minimal medium plates at pH=7.2 and exposed to the various light conditions. Homogeneous light was supplied by LED panels. Temperature was maintained at 25°C at the level of plates.

## Acknowledgments

We thank Dr. Setsuko Wakao for assistance in growing cells, Jacob Irby for performing immunodetection, and Dr. Guillaume Allorent and Dr. Giovanni Finazzi for providing the *npq4stt7* strain. A.B. acknowledges the support of the Carnegie Institution for Science. This work was supported by the U.S. Department of Energy, Office of Science, Basic Energy Sciences, Chemical Sciences, Geosciences, and Biosciences Division under the field work proposal 449B. K.K.N. is an investigator of the Howard Hughes Medical Institute.

## Competing interests

The authors declare that they have no competing interest.

## Data and materials availability

All data needed to evaluate the conclusions in the paper are present in the paper and/or the Supplementary Materials.

## Supporting Materials

77K emission spectra PAM replicates for WT and *npq4lhcsr1* (SI Fig 1)

77K emission spectra for *stt7* and *stt7npq4* (SI Fig 2)

Maximum quenching envelopes for WT and *stt7* (SI Fig 3).

Kinetics of qT in WT and *npq4lhcsr1* (SI Fig 4).

Quantification of protein contributions: distributions, averages, errors (SI Fig 5-7).

Growth of cells under constant LL or HL (SI Fig 8).

Fv/Fm values for all strains (SI Fig 9)

Kinetics of CCM-related decrease in WT NPQ during HL (SI Fig 10)

Comparison of integration results for WT and pH-sensing mutant in *Chlamydomonas* and

*Arabidopsis* (SI Fig 11)

Immunodetection of LHCSR proteins (SI Fig 12)

Quenching trajectories with error (standard deviation) for WT and *npq4lhcsr1* (SI Fig 13)

Quenching trajectories with error (standard deviation) for *lhcsr1* and *npq4* (SI Fig 14)

Quenching trajectories with error (standard deviation) for *stt7* and *stt7npq4* (SI Fig 15)

## Notes

### Competing Interest Statement

The authors have declared no competing interest.

### Summary of Updates

To fix a conversion problem on Figure 7

## References

Allen JF (1992) Protein phosphorylation in regulation of photosynthesis. Biochim. Biophys. Acta Bioenerg. 1098: 275–335 https://doi.org/10.1016/S0005-2728(09)91014-3

Allorent G, Tokutsu R, Roach T, Peers G, Cardol P, Girard-Bascou J, Seigneurin-Berny D, Petroutsos D, Kuntz M, Breyton C, Franck F, Wollman FA, Niyogi KK, Krieger-Liszkay A, Minagawa J, Finazzi G (2013) A dual strategy to cope with high light in Chlamydomonas reinhardtii. Plant Cell 25: 545–557 10.1105/tpc.112.108274

Amarnath K, Zaks J, Park SD, Niyogi KK, Fleming GR (2012) Fluorescence lifetime snapshots reveal two rapidly reversible mechanisms of photoprotection in live cells of Chlamydomonas reinhardtii. Proc. Natl. Acad. Sci. U.S.A 109: 8405–8410 10.1073/pnas.1205303109

Aro E-M, Virgin I, Andersson B (1993) Photoinhibition of photosystem II. inactivation, protein damage and turnover. Biochim. Biophys. Acta Bioenerg. 1143: 113–134 https://doi.org/10.1016/0005-2728(93)90134-2

Ballottari M, Truong TB, De Re E, Erickson E, Stella GR, Fleming GR, Bassi R, Niyogi KK (2016) Identification of pH-sensing Sites in the Light Harvesting Complex Stress-related 3 Protein Essential for Triggering Non-photochemical Quenching in Chlamydomonas reinhardtii. J. Biol. Chem. 291: 7334–7346 10.1074/jbc.M115.704601

Barera S, Dall’Osto L, Bassi R (2021) Effect of lhcsr gene dosage on oxidative stress and light use efficiency by Chlamydomonas reinhardtii cultures. J. Biotechnol. 328: 12–22 https://doi.org/10.1016/j.jbiotec.2020.12.023

Bellafiore S, Barneche F, Peltier G, Rochaix J-D (2005) State transitions and light adaptation require chloroplast thylakoid protein kinase STN7. Nature 433: 892–895 10.1038/nature03286

Benedetti M, Vecchi V, Barera S, Dall’Osto L (2018) Biomass from microalgae: the potential of domestication towards sustainable biofactories. Microb. Cell Factory 17: 173 10.1186/s12934-018-1019-3

Bergner SV, Scholz M, Trompelt K, Barth J, Gäbelein P, Steinbeck J, Xue H, Clowez S, Fucile G, Goldschmidt-Clermont M, Fufezan C, Hippler M (2015) STATE TRANSITION7-Dependent phosphorylation is modulated by changing environmental conditions, and its absence triggers remodeling of photosynthetic protein complexes. Plant Physiol. 168: 615–634 10.1104/pp.15.00072

Björkman O, Demmig B (1987) Photon yield of O<sub>2</sub> evolution and chlorophyll fluorescence characteristics at 77 K among vascular plants of diverse origins. Plantae 170: 489–504 10.1007/BF00402983

Bonaventura C, Myers J (1969) Fluorescence and oxygen evolution from Chlorella pyrenoidosa. Biochim. Biophys. Acta Bioenerg. 189: 366–383 https://doi.org/10.1016/0005-2728(69)90168-6

Bonente G, Ballottari M, Truong TB, Morosinotto T, Ahn TK, Fleming GR, Niyogi KK, Bassi R (2011) Analysis of LhcSR3, a protein essential for feedback de-excitation in the green alga Chlamydomonas reinhardtii. Plos Biol. 9: e1000577 10.1371/journal.pbio.1000577

Briantais JM, Vernotte C, Picaud M, Krause GH (1979) A quantitative study of the slow decline of chlorophyll a fluorescence in isolated chloroplasts. Biochim. Biophys. Acta Bioenerg. 548: 128–138 https://doi.org/10.1016/0005-2728(79)90193-2

Bru P, Steen CJ, Park S, Amstutz CL, Sylak-Glassman EJ, Leuenberger M, Lam L, Longoni F, Fleming GR, Niyogi KK, Malnoë A (2021) Photoprotective qH occurs in the light-harvesting complex II trimer. BioRxiv: 2021.2007.2009.450705 10.1101/2021.07.09.450705

Burlacot A, Dao O, Auroy P, Cuiné S, Li-Beisson Y, Peltier G (2021) Alternative electron pathways of photosynthesis drive the algal CO<sub>2</sub> concentrating mechanism. BioRxiv: 2021.2002.2025 432959 10.1101/2021.02.25.432959

Cantrell M, Peers G (2017) A mutant of Chlamydomonas without LHCSR maintains high rates of photosynthesis, but has reduced cell division rates in sinusoidal light conditions. Plos One 12: e0179395 10.1371/journal.pone.0179395

Cardol P, Alric J, Girard-Bascou J, Franck F, Wollman FA, Finazzi G (2009) Impaired respiration discloses the physiological significance of state transitions in Chlamydomonas. 106: 15979–15984 10.1073/pnas.0908111106

Casper-Lindley C, Björkman O (1996) Nigericin insensitive post-illumination reduction in fluorescence yield in Dunaliella tertiolecta (chlorophyte). Photosynth. Res. 50: 209–222 10.1007/BF00033120

Correa-Galvis V, Redekop P, Guan K, Griess A, Truong TB, Wakao S, Niyogi KK, Jahns P (2016) Photosystem II subunit PsbS is involved in the induction of LHCSR protein-dependent energy dissipation in Chlamydomonas reinhardtii. J. Biol. Chem. 291: 17478–17487 10.1074/jbc.M116.737312

Dall’Osto L, Caffarri S, Bassi R (2005) A Mechanism of Nonphotochemical Energy Dissipation, Independent from PsbS, Revealed by a Conformational Change in the Antenna Protein CP26. Plant Cell 17: 1217–1232 10.1105/tpc.104.030601

Depege N, Bellafiore S, Rochaix JD (2003) Role of chloroplast protein kinase Stt7 in LHCII phosphorylation and state transition in Chlamydomonas. Science 299: 1572–1575

Dinc E, Tian L, Roy LM, Roth R, Goodenough U, Croce R (2016) LHCSR1 induces a fast and reversible pH-dependent fluorescence quenching in LHCII in Chlamydomonas reinhardtii cells. Proc. Nat. Acad. Sci. U. S. A. 113: 7673–7678 10.1073/pnas.1605380113

Drop B, Webber-Birungi M, Yadav SKN, Filipowicz-Szymanska A, Fusetti F, Boekema EJ, Croce R (2014) Light-harvesting complex II (LHCII) and its supramolecular organization in Chlamydomonas reinhardtii. Biochim. Biophys. Acta Bioenerg. 1837: 63–72 https://doi.org/10.1016/j.bbabio.2013.07.012

Erickson E, Wakao S, Niyogi KK (2015) Light stress and photoprotection in Chlamydomonas reinhardtii. Plant J. 82: 449–465 10.1111/tpj.12825

Fei C, Wilson AT, Mangan NM, Wingreen NS, Jonikas MC (2021) Diffusion barriers and adaptive carbon uptake strategies enhance the modeled performance of the algal CO<sub>2</sub> concentrating mechanism. BioRxiv: 2021.2003.2004.433933 10.1101/2021.03.04.433933

Forti G, Caldiroli G (2005) State transitions in Chlamydomonas reinhardtii. The role of the mehler reaction in state 2-to-state 1 transition. Plant Physiol. 137: 492–499 10.1104/pp.104.048256

Gao S, Pinnola A, Zhou L, Zheng Z, Li Z, Bassi R, Wang G (2021) Light-harvesting complex stress-related proteins play crucial roles in the acclimation of Physcomitrella patens under fluctuating light conditions. Phot. Res 10.1007/s11120-021-00874-8

Girolomoni L, Cazzaniga S, Pinnola A, Perozeni F, Ballottari M, Bassi R (2019) LHCSR3 is a nonphotochemical quencher of both photosystems in Chlamydomonas reinhardtii. Proc. Nat. Acad. Sci. U. S. A. 116: 4212–4217 10.1073/pnas.1809812116

Goral TK, Johnson MP, Duffy CDP, Brain APR, Ruban AV, Mullineaux CW (2012) Light-harvesting antenna composition controls the macrostructure and dynamics of thylakoid membranes in Arabidopsis. Plant J. 69: 289–301 https://doi.org/10.1111/j.1365-313X.2011.04790.x

Graham PJ, Nguyen B, Burdyny T, Sinton D (2017) A penalty on photosynthetic growth in fluctuating light. Scientific Rep. 7: 12513 10.1038/s41598-017-12923-1

Huang Z, Shen L, Wang W, Mao Z, Yi X, Kuang T, Shen J-R, Zhang X, Han G (2021) Structure of photosystem I-LHCI-LHCII from the green alga Chlamydomonas reinhardtii in State 2. Nature Commun. 12: 1100 10.1038/s41467-021-21362-6

Iwai M, Takizawa K, Tokutsu R, Okamuro A, Takahashi Y, Minagawa J (2010a) Isolation of the elusive supercomplex that drives cyclic electron flow in photosynthesis. Nature 464: 1210–U1134 10.1038/nature08885

Iwai M, Yokono M, Inada N, Minagawa J (2010b) Live-cell imaging of photosystem II antenna dissociation during state transitions. Proc. Natl. Acad. Sci. USA 107: 2337–2342 10.1073/pnas.0908808107

Kawakami K, Tokutsu R, Kim E, Minagawa J (2019) Four distinct trimeric forms of light-harvesting complex II isolated from the green alga Chlamydomonas reinhardtii. Phot. Res. 142: 195–201 10.1007/s11120-019-00669-y

Khorobrykh S, Havurinne V, Mattila H, Tyystjärvi E (2020) Oxygen and ROS in photosynthesis. Plants 9: 91

Klughammer C, Schreiber U (2008) Complementary PS II quantum yields calculated from simple fluorescence parameters measured by PAM fluorometry and the Saturation Pulse method. PAM application notes 1: 201–247

Kondo T, Pinnola A, Chen WJ, Dall’Osto L, Bassi R, Schlau-Cohen GS (2017) Single-molecule spectroscopy of LHCSR1 protein dynamics identifies two distinct states responsible for multi-timescale photosynthetic photoprotection. Nature Chem. 9: 772–778 10.1038/nchem.2818

Kosuge K, Tokutsu R, Kim E, Akimoto S, Yokono M, Ueno Y, Minagawa J (2018) LHCSR1-dependent fluorescence quenching is mediated by excitation energy transfer from LHCII to photosystem I in Chlamydomonas reinhardtii. Proc. Natl. Acad. Sci. USA 115: 3722–3727 10.1073/pnas.1720574115

Kromdijk J, Głowacka K, Leonelli L, Gabilly ST, Iwai M, Niyogi KK, Long SP (2016) Improving photosynthesis and crop productivity by accelerating recovery from photoprotection. Science 354: 857–861 10.1126/science.aai8878

Lemeille S, Willig A, Depège-Fargeix N, Delessert C, Bassi R, Rochaix J-D (2009) Analysis of the Chloroplast Protein Kinase Stt7 during State Transitions. Plos Biol. 7: e1000045 10.1371/journal.pbio.1000045

Liguori N, Roy LM, Opacic M, Durand G, Croce R (2013) Regulation of light harvesting in the green alga Chlamydomonas reinhardtii: the C-terminus of LHCSR Is the knob of a dimmer switch. J. Am. Chem. Soc. 135: 18339–18342 10.1021/ja4107463

Malnoë A, Schultink A, Shahrasbi S, Rumeau D, Havaux M, Niyogi KK (2018) The Plastid Lipocalin LCNP Is Required for Sustained Photoprotective Energy Dissipation in Arabidopsis. Plant Cell 30: 196–208 10.1105/tpc.17.00536

Minagawa J (2011) State transitions—The molecular remodeling of photosynthetic supercomplexes that controls energy flow in the chloroplast. Biochim. Biophys. Acta Bioenerg. 1807: 897–905 https://doi.org/10.1016/j.bbabio.2010.11.005

Nagy G, Ünnep R, Zsiros O, Tokutsu R, Takizawa K, Porcar L, Moyet L, Petroutsos D, Garab G, Finazzi G, Minagawa J (2014) Chloroplast remodeling during state transitions in Chlamydomonas reinhardtii as revealed by noninvasive techniques in vivo. Proc. Natl. Acad. Sci. U.S.A 111: 5042 10.1073/pnas.1322494111

Nawrocki WJ, Liu X, Croce R (2020) Chlamydomonas reinhardtii exhibits de facto constitutive NPQ capacity in physiologically relevant conditions. Plant Physiol. 182: 472–479 10.1104/pp.19.00658

Nawrocki WJ, Liu X, Raber B, Hu C, de Vitry C, Bennett DIG, Croce R (2021) Molecular origins of induction and loss of photoinhibition-related energy dissipation qI. Sci. Adv. 7: 2021.2003.2010.434601 10.1126/sciadv.abj0055

Nawrocki WJ, Santabarbara S, Mosebach L, Wollman F-A, Rappaport F (2016) State transitions redistribute rather than dissipate energy between the two photosystems in Chlamydomonas. Nat. Plant 2: 16031 10.1038/nplants.2016.31

Nedbal L, Lazár D (2021) Photosynthesis dynamics and regulation sensed in the frequency domain. Plant Physiol. 187: 646–661 10.1093/plphys/kiab317

Nilkens M, Kress E, Lambrev P, Miloslavina Y, Müller M, Holzwarth AR, Jahns P (2010) Identification of a slowly inducible zeaxanthin-dependent component of non-photochemical quenching of chlorophyll fluorescence generated under steady-state conditions in Arabidopsis. Biochim. Biophys. Acta Bioenerg. 1797: 466–475 https://doi.org/10.1016/j.bbabio.2010.01.001

Niyogi KK, Bjorkman O, Grossman AR (1997) Chlamydomonas Xanthophyll Cycle Mutants Identified by Video Imaging of Chlorophyll Fluorescence Quenching. Plant Cell 9: 1369–1380 10.1105/tpc.9.8.1369

Peers G, Truong TB, Ostendorf E, Busch A, Elrad D, Grossman AR, Hippler M, Niyogi KK (2009) An ancient light-harvesting protein is critical for the regulation of algal photosynthesis. Nature 462: 518–521

Perin G, Jones PR (2019) Economic feasibility and long-term sustainability criteria on the path to enable a transition from fossil fuels to biofuels. Curr. Opin. Biotechnol. 57: 175–182 https://doi.org/10.1016/j.copbio.2019.04.004

Perozeni F, Beghini G, Cazzaniga S, Ballottari M (2020) Chlamydomonas reinhardtii LHCSR1 and LHCSR3 proteins involved in photoprotective non-photochemical quenching have different quenching efficiency and different carotenoid affinity. Sci. Rep. 10: 21957 10.1038/s41598-020-78985-w

Pinnola A, Bassi R (2018) Molecular mechanisms involved in plant photoprotection. Biochem. Soc. Trans. 46: 467–482 10.1042/bst20170307

Rintamäki E, Martinsuo P, Pursiheimo S, Aro E-M (2000) Cooperative regulation of light-harvesting complex II phosphorylation via the plastoquinol and ferredoxin-thioredoxin system in chloroplasts. Proc. Natl. Acad. Sci. USA 97: 11644–11649 doi:10.1073/pnas.180054297

Roach T (2020) LHCSR3-Type NPQ Prevents Photoinhibition and Slowed Growth under Fluctuating Light in Chlamydomonas reinhardtii. Plants 9: 1604

Roach T, Na CS (2017) LHCSR3 affects de-coupling and re-coupling of LHCII to PSII during state transitions in Chlamydomonas reinhardtii. Sci. Rep. 7: 43145 10.1038/srep43145

Roach T, Na CS, Stöggl W, Krieger-Liszkay A (2020) The non-photochemical quenching protein LHCSR3 prevents oxygen-dependent photoinhibition in Chlamydomonas reinhardtii. J. Exp. Botany 71: 2650–2660 10.1093/jxb/eraa022

Rochaix J-D, Bassi R (2019) LHC-like proteins involved in stress responses and biogenesis/repair of the photosynthetic apparatus. Biochem. J. 476: 581–593 10.1042/BCJ20180718

Semchonok DA, Sathish Yadav KN, Xu P, Drop B, Croce R, Boekema EJ (2017) Interaction between the photoprotective protein LHCSR3 and C2S2 Photosystem II supercomplex in Chlamydomonas reinhardtii. Biochim. Biophys. Acta Bioenerg. 1858: 379–385 https://doi.org/10.1016/j.bbabio.2017.02.015

Steen CJ, Morris JM, Short AH, Niyogi KK, Fleming GR (2020) Complex Roles of PsbS and Xanthophylls in the Regulation of Nonphotochemical Quenching in Arabidopsis thaliana under Fluctuating Light. J. Phys. Chem. B 124: 10311–10325 10.1021/acs.jpcb.0c06265

Sylak-Glassman EJ, Malnoë A, De Re E, Brooks MD, Fischer AL, Niyogi KK, Fleming GR (2014) Distinct roles of the photosystem II protein PsbS and zeaxanthin in the regulation of light harvesting in plants revealed by fluorescence lifetime snapshots. Proc. Natl. Acad. Sci. USA 111: 17498–17503 10.1073/pnas.1418317111

Sylak-Glassman EJ, Zaks J, Amarnath K, Leuenberger M, Fleming GR (2016) Characterizing non-photochemical quenching in leaves through fluorescence lifetime snapshots. Phot. Res. 127: 69–76 10.1007/s11120-015-0104-2

Tanaka Y, Adachi S, Yamori W (2019) Natural genetic variation of the photosynthetic induction response to fluctuating light environment. Curr Opin. Plant Biol. 49: 52–59 https://doi.org/10.1016/j.pbi.2019.04.010

Tian L, Dinc E, Croce R (2015) LHCII populations in different quenching states are present in the thylakoid membranes in a ratio that depends on the light conditions. J. Phys. Chem. Letters 6: 2339–2344 10.1021/acs.jpclett.5b00944

Tian L, Nawrocki WJ, Liu X, Polukhina I, van Stokkum IHM, Croce R (2019) pH dependence, kinetics and light-harvesting regulation of nonphotochemical quenching in Chlamydomonas. Proc Nat. Acad. Sci. USA 116: 8320–8325 10.1073/pnas.1817796116

Tibiletti T, Auroy P, Peltier G, Caffarri S (2016) Chlamydomonas reinhardtii PsbS protein is functional and accumulates rapidly and transiently under high light. Plant Physiol. 171: 2717–2730 10.1104/pp.16.00572

Troiano JM, Perozeni F, Moya R, Zuliani L, Baek K, Jin E, Cazzaniga S, Ballottari M, Schlau-Cohen GS (2021) Identification of distinct pH-and zeaxanthin-dependent quenching in LHCSR3 from Chlamydomonas reinhardtii. elife 10: e60383 10.7554/eLife.60383

Truong TB (2011) Investigating the role(s) of LHCSRs in Chlamydomonas reinhardtii. University of California, Berkeley

Ünlü C, Drop B, Croce R, van Amerongen H (2014) State transitions in Chlamydomonas reinhardtii strongly modulate the functional size of photosystem II but not of photosystem I. Proc. Natl. Acad. Sci. U.S.A 111: 3460–3465 10.1073/pnas.1319164111

Vecchi V, Barera S, Bassi R, Dall’Osto L (2020) Potential and challenges of improving photosynthesis in algae. Plants 9: 67

Vink M, Zer H, Alumot N, Gaathon A, Niyogi K, Herrmann RG, Andersson B, Ohad I (2004) Light-modulated exposure of the light-harvesting complex II (LHCII) to protein kinase(s) and state transition in Chlamydomonas reinhardtii xanthophyll mutants. Biochem. 43: 7824–7833 10.1021/bi030267l

Wang Y, Burgess SJ, de Becker EM, Long SP (2020) Photosynthesis in the fleeting shadows: an overlooked opportunity for increasing crop productivity? Plant J. 101: 874–884 https://doi.org/10.1111/tpj.14663

Wehner A, Grasses T, Jahns P (2006) De-epoxidation of Violaxanthin in the Minor Antenna Proteins of Photosystem II, LHCB4, LHCB5, and LHCB6 J. Biol. Chem. 281: 21924–21933 10.1074/jbc.M602915200

Zaks J, Amarnath K, Kramer DM, Niyogi KK, Fleming GR (2012) A kinetic model of rapidly reversible nonphotochemical quenching. Proc. Nat. Acad. U.S.A 109: 15757–15762 10.1073/pnas.1211017109

Zaks J, Amarnath K, Sylak-Glassman EJ, Fleming GR (2013) Models and measurements of energy-dependent quenching. Photosynth. Res. 116: 389–409 10.1007/s11120-013-9857-7

Zhang XJ, Fujita Y, Tokutsu R, Minagawa J, Ye S, Shibata Y (2021) High-Speed Excitation-Spectral Microscopy Uncovers In Situ Rearrangement of Light-Harvesting Apparatus in Chlamydomonas during State Transitions at Submicron Precision. Plant Cell Physiol. 62: 872–882 10.1093/pcp/pcab047

