## Supplemental informations for "Interplay between LHCSR proteins and state transitions governs the NPQ response in intact cells of *Chlamydomonas* during light fluctuations"

### for

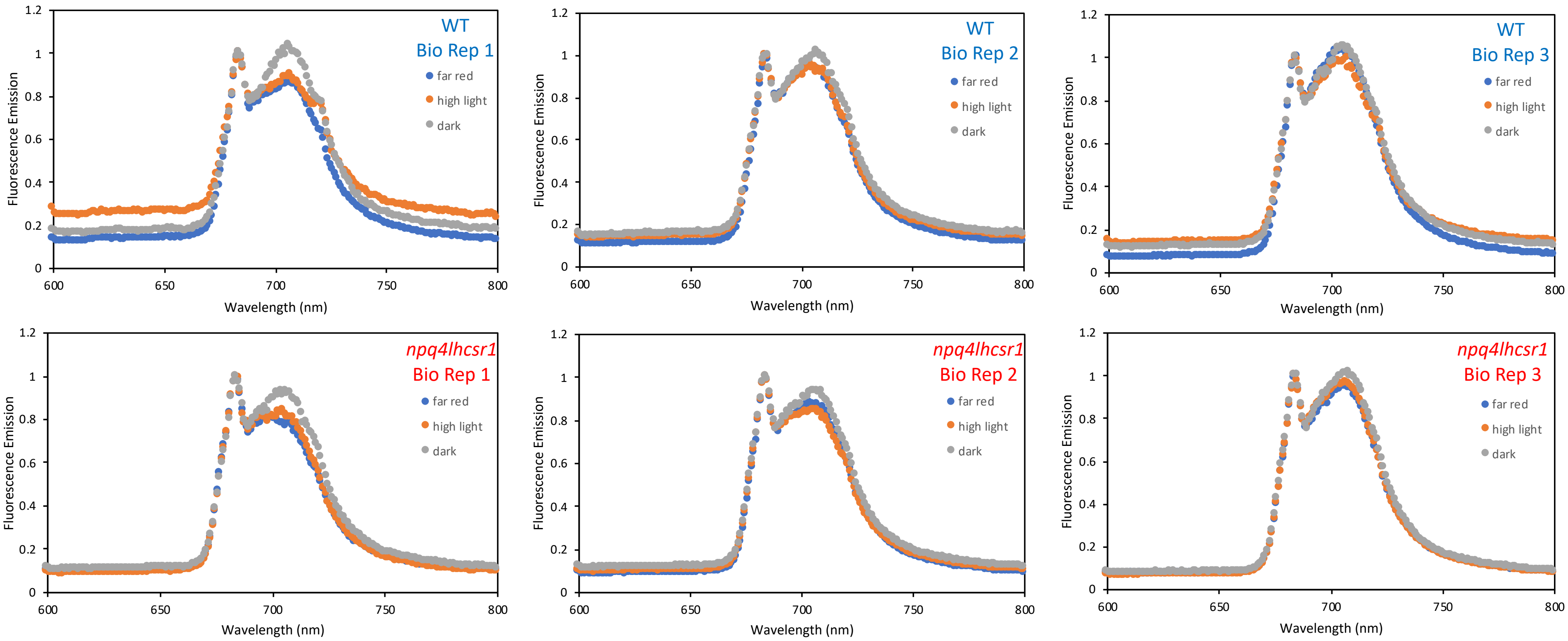


**Supporting Figure 1**: Independent biological replicates of 77K fluorescence emission spectra for WT and *npq4lhcsr1*. Within each graph, the spectra colored in blue, orange, and gray represent those taken at 0 min (far-red acclimated), 10 min (HL-exposed cells), and 20 min (dark-exposed cells) timepoints, respectively. Samples were taken under PAM conditions. [*supports Fig. 4 of main text*]

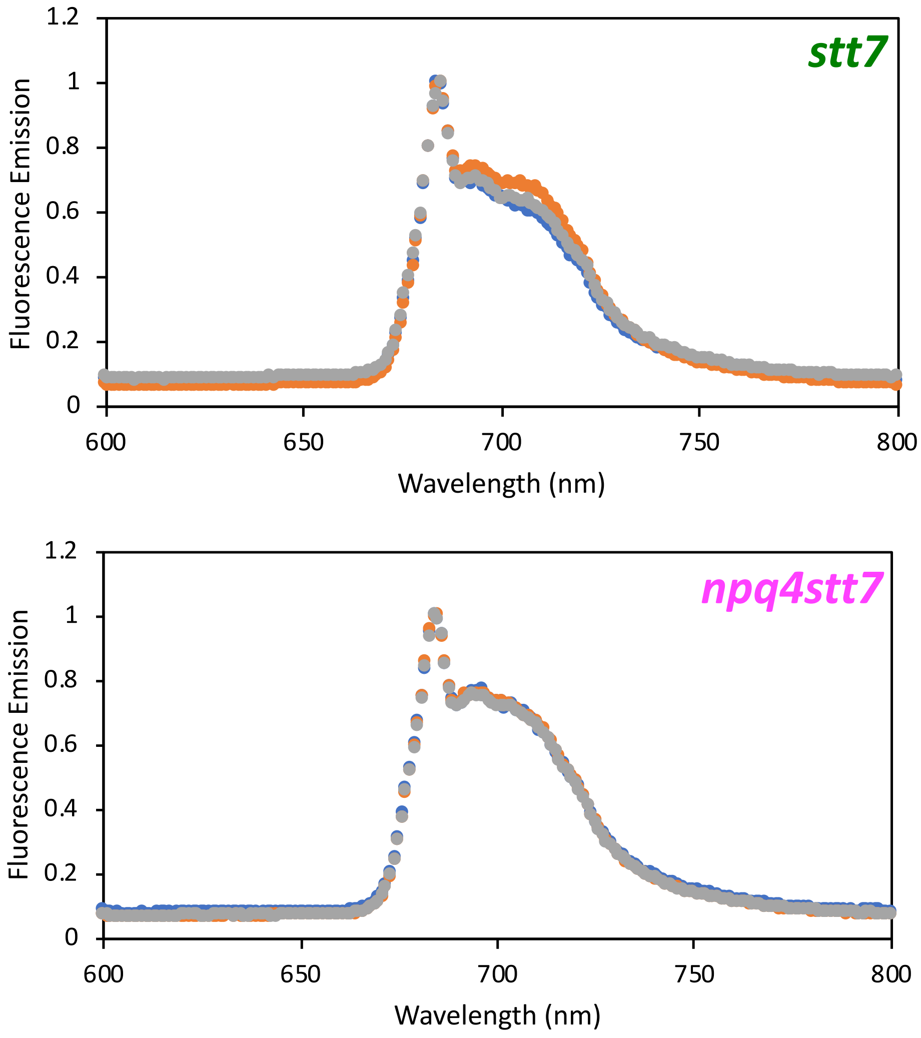


**Supporting Figure 2**: Representative 77K fluorescence emission spectra for *stt7* and *npq4stt7* strains. For each strain, the spectra colored in blue, orange, and gray represent those taken at 0 min (far-red acclimated), 10 min (HL-exposed cells), and 20 min (dark-exposed cells) timepoints, respectively. Samples were taken under TCSPC conditions. [*supports Fig. 4 of main text*]

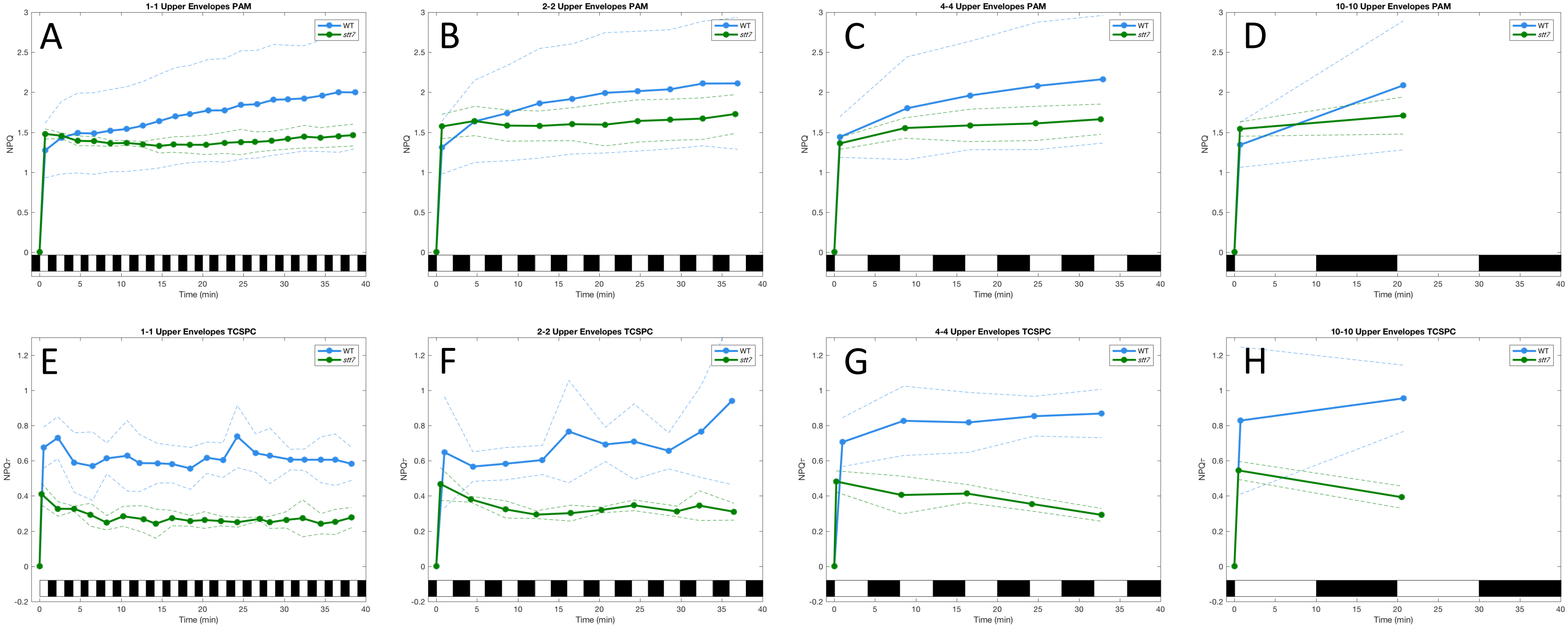


**Supporting Figure 3.** Maximum quenching envelopes defined by the maximum values of NPQ (measured by PAM, **A-D**) or NPQτ (measured by TCSPC, **E-H**) during every high light period for WT (blue) and *stt7* (green). Solid lines denote the experimental data points, while dashed lines above and below the curves show the addition and subtraction of the experimental standard deviation. Note that when comparing the envelopes between the two measurement techniques, there are qualitative differences in the envelope structure. In the case of the PAM measurements, WT shows a maximum NPQ value that gradually increases as light fluctuations progress (due to activation of qT as the experiment progresses), while *stt7* shows no such increase (due to the absence of qT). For TCSPC measurements, WT shows a maximum NPQτ value that is relatively flat as light fluctuations progress (includes the effects of qT), while *stt7* shows a slightly decreasing quenching envelope (due to the absence of qT).

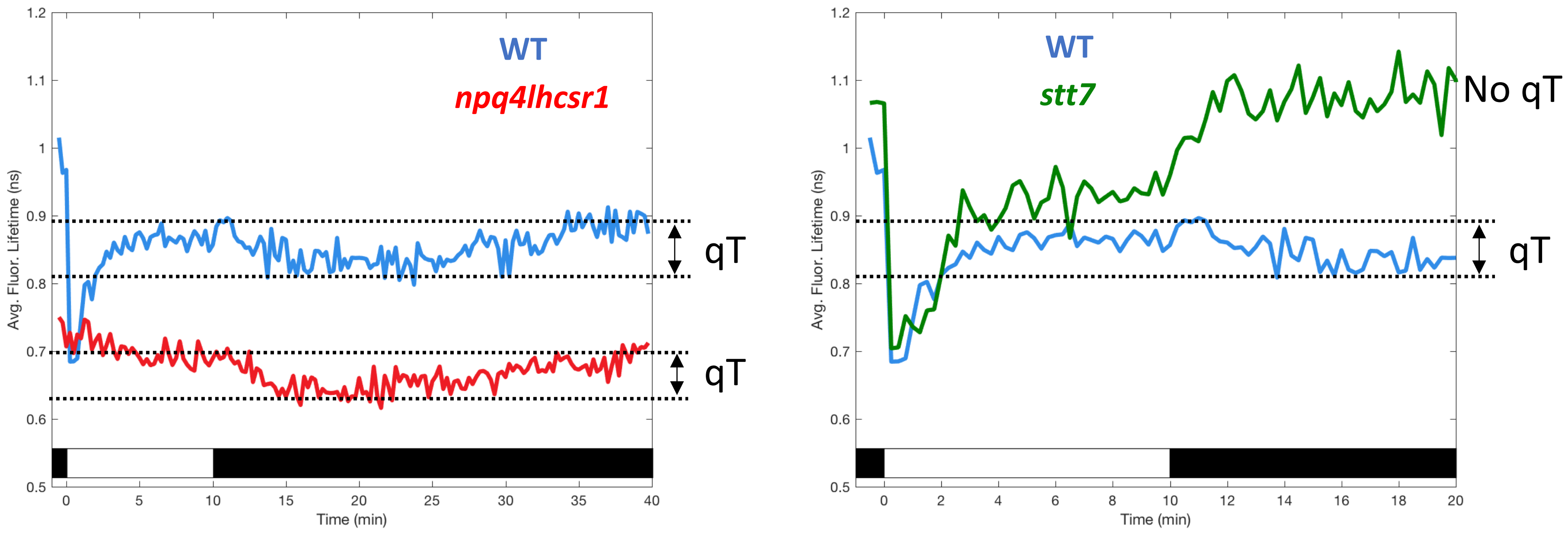


**Supporting Figure 4.** Kinetics of qT measured by TCSPC. Following 15 min of far-red acclimation, cells were exposed to 10 min of HL followed by 30 min of darkness (left panel) or 10 min HL followed by 10 min of darkness (right panel). WT is shown in blue, *npq4lhcsr1* in red, and *stt7* in green. Each lifetime trajectory is the average of 3 biological replicates where each biological replicate is average of 3 technical replicates. Dashed horizontal lines highlight the extent of the decreasing lifetime observed upon HL-to-dark transition in STT7-containing lines. Note that after 10 to 15 min of quenching (evidenced by the decreasing average fluorescence lifetimes in each strain), the quenching then begins to turn off and eventually returns to the starting lifetime. The *stt7* mutant shows no such decrease in lifetime during a 10-minute dark period.

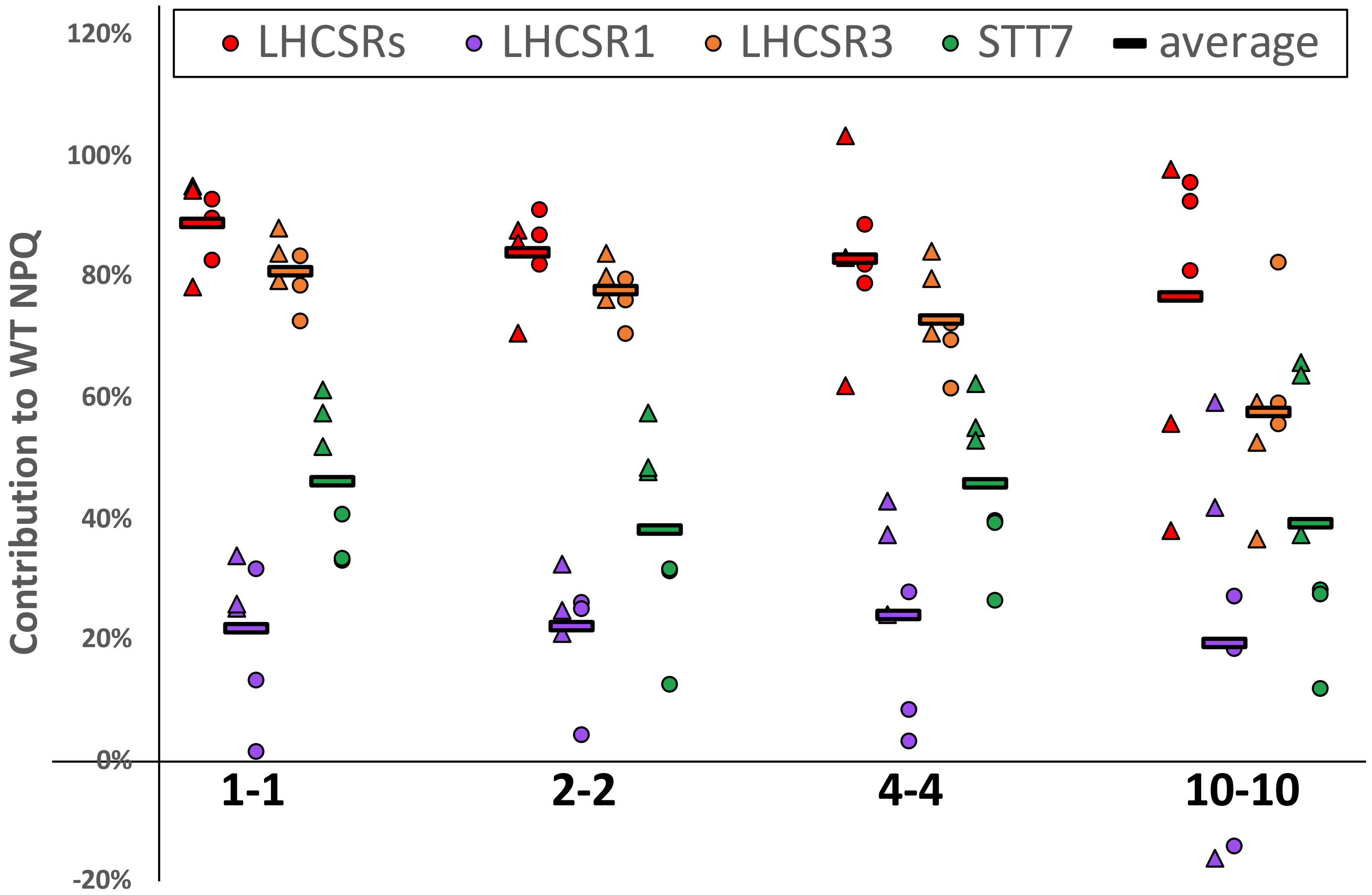


**Supporting Figure 5.** Distribution of replicate measurements for the contribution of each protein to overall WT NPQ as a function of fluctuating light sequence. Each data point represents the percentage of the WT NPQ that is lost in each mutant following integration of the trajectories of NPQ (measured by PAM, *circles*) or NPQτ (measured by TCSPC, *triangles*). The contribution of each protein is colored according to the respective mutant that was used for quantification relative to WT (*red,* LHCSRs from *npq4lhcsr1*; *purple*, LHCSR1 from *lhcsr1*; *orange*, LHCSR3 from *npq4*; *green*, STT7 from *stt7*). Within each colored cluster, the triangles represent the 3 biological replicates measured by TCSPC (each biological replicate is the average of 3 technical replicates) and the circles represent the 3 biological replicates measured by PAM. The colored horizontal lines represent the average of all TCSPC and PAM data points for each protein.

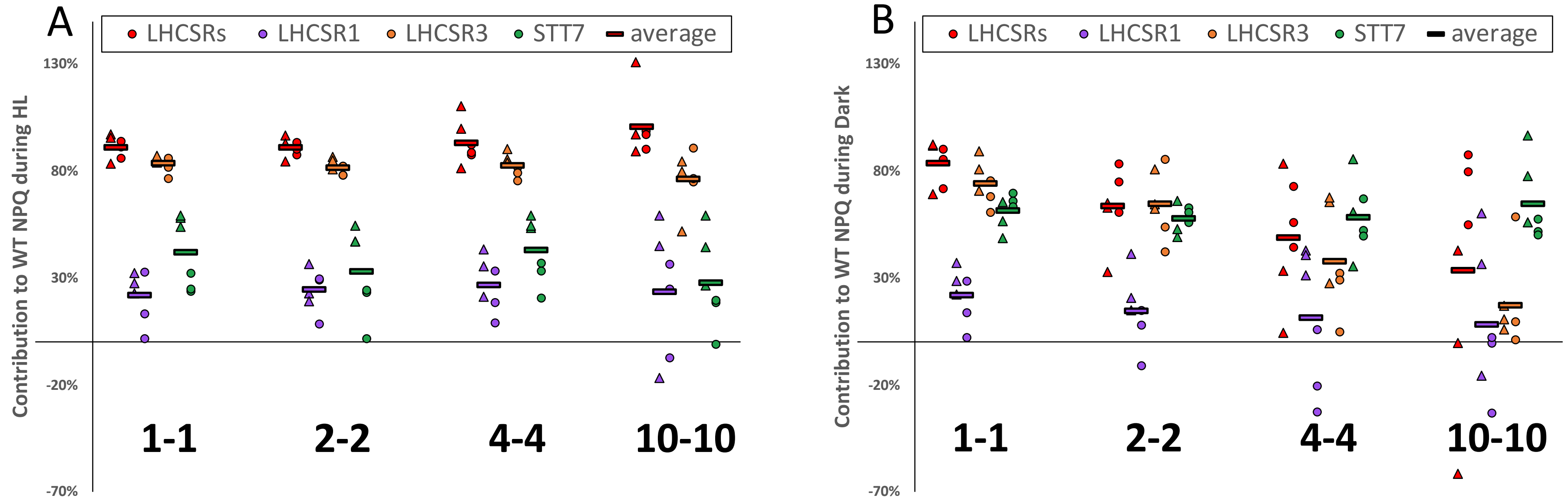


| HL Contributions | 1-1 | 2-2 | 4-4 | 10-10 | AVERAGE |
|---|---|---|---|---|---|
| LHCSRs | 91 ± 5% | 91 ± 4% | 93 ± 10% | 100 ± 15% | 94% |
| LHCSR1 | 22 ± 12% | 24 ± 10% | 27 ± 13% | 23 ± 30% | 24% |
| LHCSR3 | 83 ± 4% | 82 ± 4% | 82 ± 5% | 74 ± 13% | 81% |
| STT7 | 42 ± 17% | 33 ± 20% | 43 ± 15% | 28 ± 21% | 36% |

| Dark Contributions | 1-1 | 2-2 | 4-4 | 10-10 | AVERAGE |
|---|---|---|---|---|---|
| LHCSRs | 83 ± 10% | 62 ± 17% | 49 ± 28% | 34 ± 56% | 57% |
| LHCSR1 | 22 ± 13% | 15 ± 17% | 11 ± 33% | 8 ± 35% | 14% |
| LHCSR3 | 74 ± 10% | 65 ± 16% | 38 ± 25% | 17 ± 21% | 48% |
| STT7 | 61 ± 7% | 58 ± 6% | 58 ± 17% | 65 ± 19% | 60% |

| Ratio HL/Dark | 1-1 | 2-2 | 4-4 | 10-10 | AVERAGE |
|---|---|---|---|---|---|
| LHCSRs | 1.09 | 1.43 | 1.91 | 2.99 | 1.86 |
| LHCSR1 | 0.99 | 1.66 | 2.41 | 2.87 | 1.98 |
| LHCSR3 | 1.13 | 1.26 | 2.19 | 4.48 | 2.27 |
| STT7 | 0.68 | 0.57 | 0.73 | 0.43 | 0.60 |

Ratio >1: larger role in HL
Ratio <1: larger role in Dark

| Difference HL -Dark | 1-1 | 2-2 | 4-4 | 10-10 | AVERAGE |
|---|---|---|---|---|---|
| LHCSRs | 7.8% | 27.5% | 44.3% | 66.8% | 37% |
| LHCSR1 | -0.3% | 9.6% | 15.6% | 15.3% | 10% |
| LHCSR3 | 9.3% | 16.9% | 44.9% | 59.2% | 33% |
| STT7 | -19.5% | -24.9% | -15.4% | -37.0% | -24% |

Positive: larger role in HL
Negative: larger role in Dark

**Supporting Figure 6.** Distribution of replicate measurements for the contribution of each protein to WT NPQ during HL **(A)** or dark (**B**) as a function of fluctuating light sequence. The tables below show the ratio of the contribution of each protein during HL to dark (obtained by dividing the two values) or the difference in contribution in HL and dark (obtained by subtracting the two values). All other details are as described in **Supp Fig. 5.**

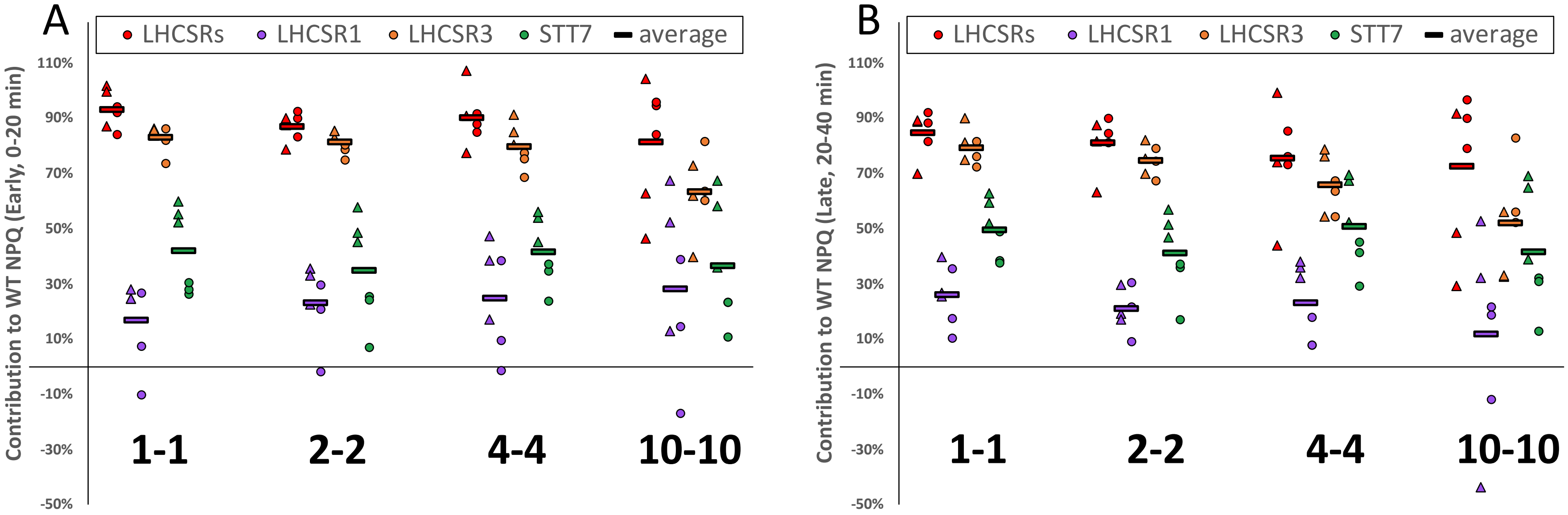


| Early Contributions | 1-1 | 2-2 | 4-4 | 10-10 | AVERAGE |
|---|---|---|---|---|---|
| LHCSRs | 93 ± 7% | 87 ± 5% | 90 ± 10% | 81 ± 22% | 88% |
| LHCSR1 | 17 ± 15% | 23 ± 14% | 25 ± 19% | 28 ± 31% | 23% |
| LHCSR3 | 83 ± 5% | 81 ± 4% | 80 ± 8% | 63 ± 14% | 77% |
| STT7 | 42 ± 16% | 35 ± 19% | 42 ± 12% | 36 ± 22% | 39% |

| Late Contributions | 1-1 | 2-2 | 4-4 | 10-10 | AVERAGE |
|---|---|---|---|---|---|
| LHCSRs | 85 ± 8% | 81 ± 9% | 75 ± 18% | 72 ± 27% | 78% |
| LHCSR1 | 26 ± 11% | 21 ± 8% | 23 ± 14% | 12 ± 34% | 20% |
| LHCSR3 | 79 ± 6% | 75 ± 6% | 66 ± 10% | 52 ± 18% | 68% |
| STT7 | 50 ± 11% | 41 ± 14% | 51 ± 16% | 41 ± 22% | 46% |

| Ratio Early/Late | 1-1 | 2-2 | 4-4 | 10-10 | AVERAGE |
|---|---|---|---|---|---|
| LHCSRs | 1.10 | 1.07 | 1.20 | 1.12 | 1.12 |
| LHCSR1 | 0.65 | 1.10 | 1.07 | 2.43 | 1.31 |
| LHCSR3 | 1.04 | 1.09 | 1.21 | 1.21 | 1.14 |
| STT7 | 0.85 | 0.85 | 0.82 | 0.88 | 0.85 |

Ratio >1: larger role early
Ratio <1: larger role later

| Difference Early-Late | 1-1 | 2-2 | 4-4 | 10-10 | AVERAGE |
|---|---|---|---|---|---|
| LHCSRs | 8.2% | 5.8% | 14.7% | 8.7% | 9% |
| LHCSR1 | -9.1% | 2.1% | 1.6% | 16.5% | 3% |
| LHCSR3 | 3.6% | 6.5% | 13.8% | 11.1% | 9% |
| STT7 | -7.7% | -6.3% | -9.0% | -5.0% | -7% |

Positive: larger role early
Negative: larger role late

**Supporting Figure 7.** Distribution of replicate measurements for the contribution of each protein to WT NPQ during the early portion (0-20 min, **A**) or during the later portion of the experiment (20-40 min, **B**) as a function of fluctuating light sequence. The tables below show the ratio of the contribution of each protein during HL to dark (obtained by dividing the two values) or the difference in contribution in HL and dark (obtained by subtracting the two values). All other details are as described in **Supp Fig. 5.**

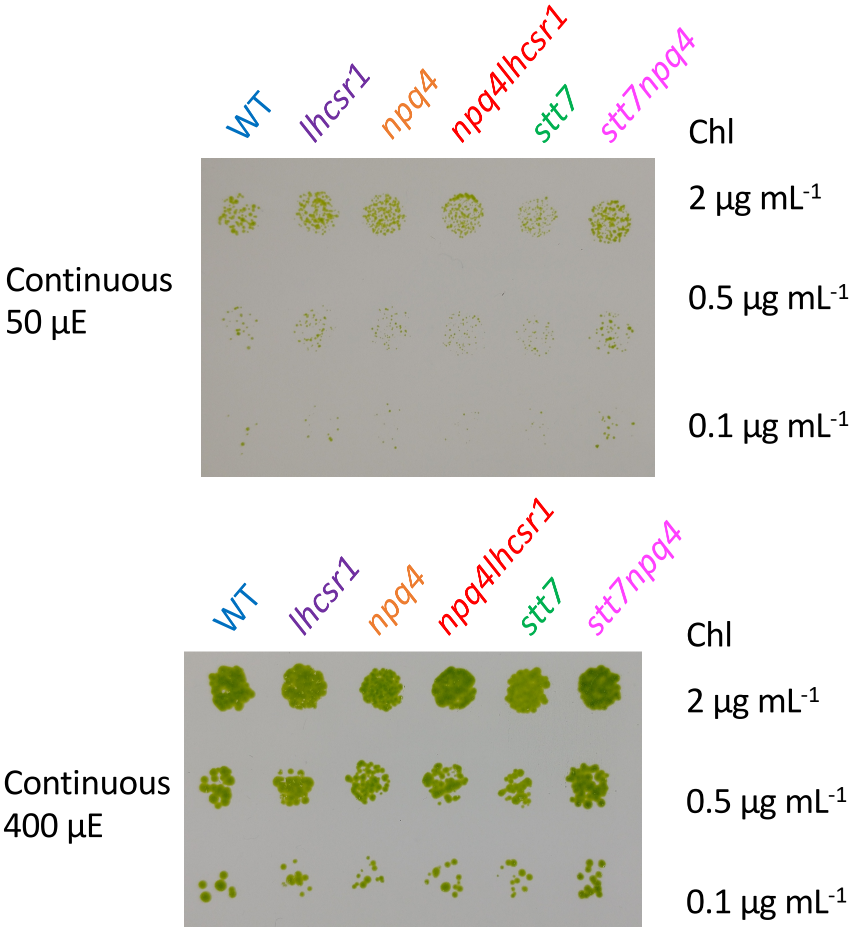


**Supporting Figure 8.** Growth of cells under continuous LL (50 uE) or HL (400 uE) conditions. All other details are as described in Fig. 7 of the main text and in the Methods. [*supports Fig. 7 of main text*]

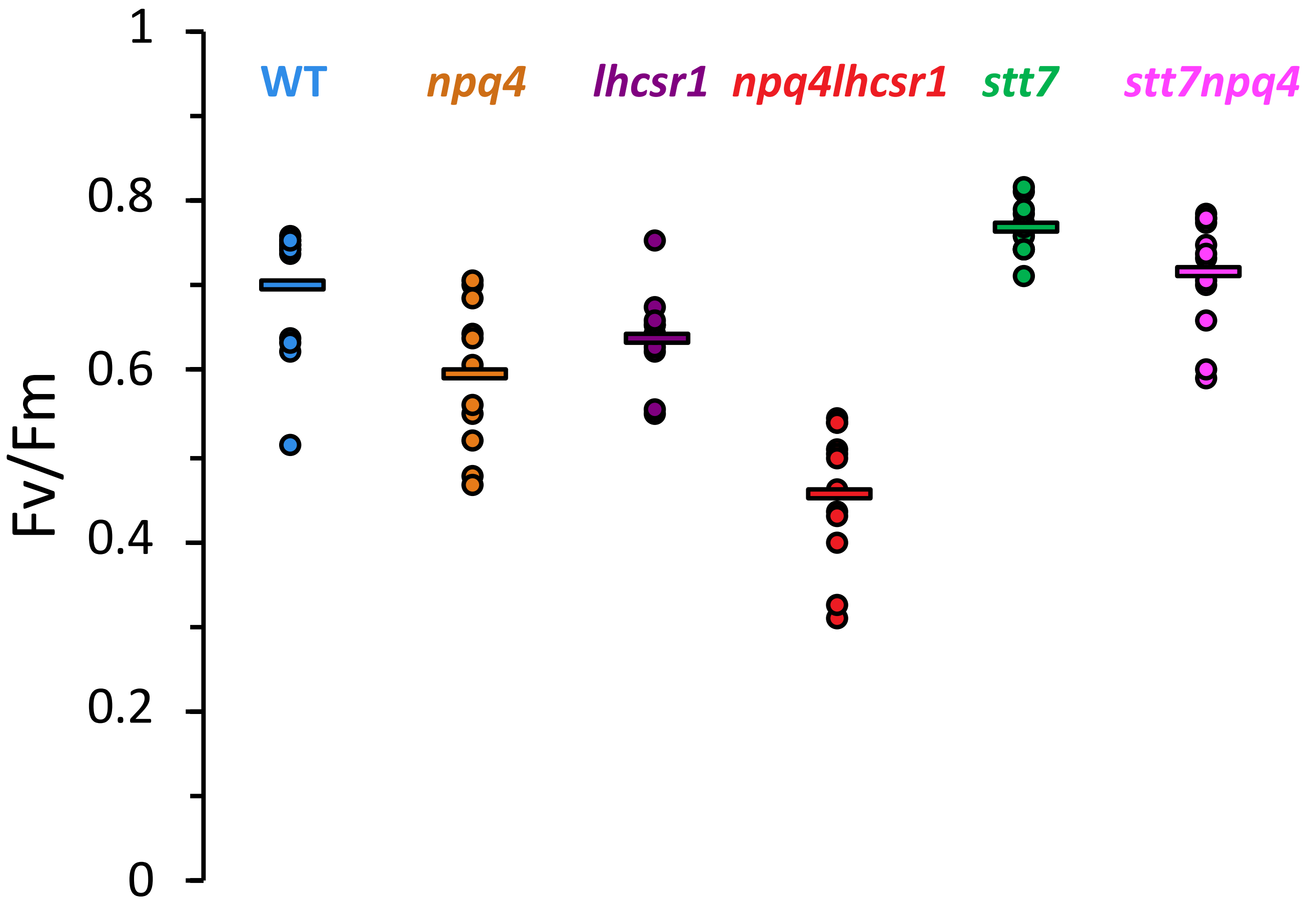


**Supporting Figure 9.** Fv/Fm values for each strain measured prior to exposure to fluctuating light sequences. Circles represent independent biological replicates. Horizontal lines show the average of 12 replicate measurements.

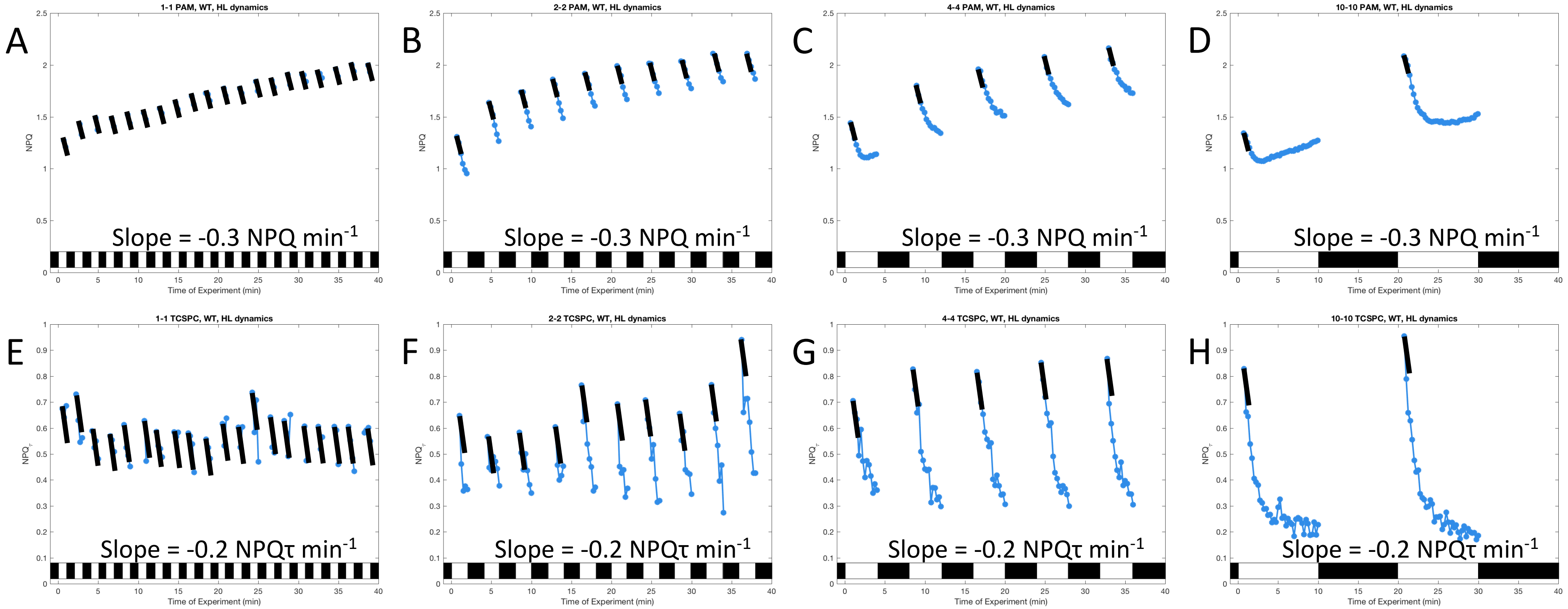


**Supporting Figure 10.** Analysis of the kinetics associated with the decrease in NPQ (measured by PAM, **A-D**) or NPQτ (measured by TCSPC, **E-H**) during every high light (HL) period for the WT strain. Within each HL period, the maximum value of NPQ (or NPQτ) and all subsequent data points after the maximum are shown (blue circles). Superimposed are solid black line segments. These lines segments have identical length and slope for panels **A-D** (PAM) and **E-H** (TCSPC). The line segments adequately capture the decrease of NPQ or NPQτ in each HL period, corresponding to approximate rates of 0.3 units of NPQ per minute (**A-D**, PAM) and 0.2 units of NPQτ per minute (**E-H**, TCSPC). As the fluctuating light period increases, a larger extent of the decrease in both NPQ and NPQτ is observed.

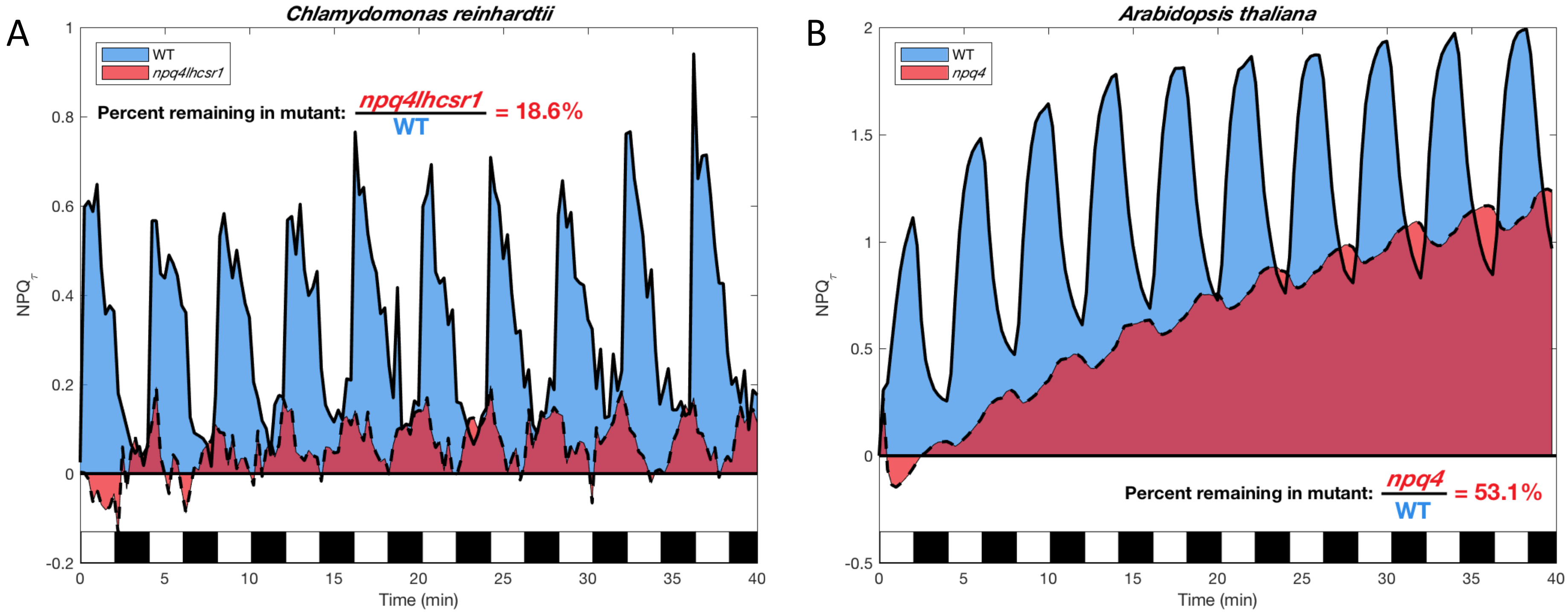


**Supporting Figure 11.** Comparison of integration results for *C. reinhardtii* cells (left) or *A. thaliana* leaves (right) exposed to 2 min HL – 2 min dark fluctuating actinic light sequence. The quantification of the amount of NPQτ remaining in each mutant lacking pH-sensing protein (*C. reinhardtii* LHCSR1 and LHCSR3; *A. thaliana* PsbS) is shown. Both were measured by TCSPC. The *C. reinhardtii* data is taken from **Fig. 2** of the main text. The *A. thaliana* data was derived from a previous paper published by our groups (Steen et al ***JPCB*** 2020).
[*supports discussion section of main text*]

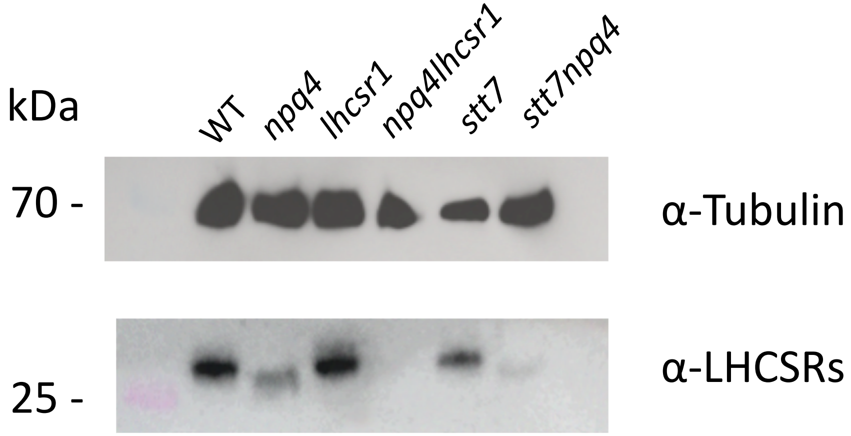


**Supporting Figure 12.** Immunodetection of LHCSRs in mutants and their control strain. Cells were harvested before the measurements described in **Fig. 1**. Immunoanalysis of LHCSRs proteins was carried out using a previously described LHCSRs antibody (Richard et al, 2000). A tubilin antibody (Agrisera XXX) was used as a loading control.

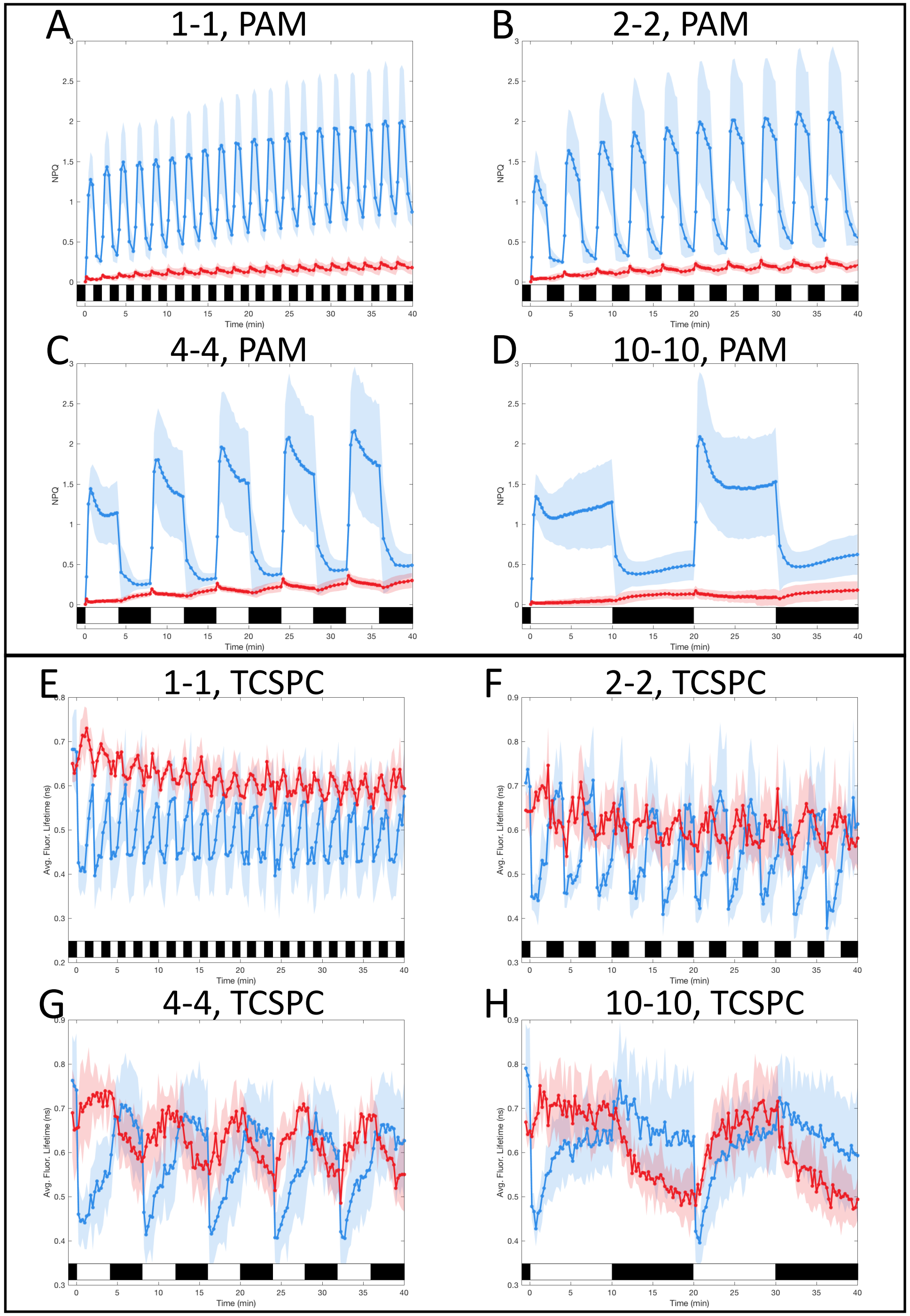


**WT**

***npq4lhcsr1***

**Supporting Figure 13.** Quenching trajectories during light fluctuations in *npq4lhcsr1* and its control strain. The response of NPQ (**A-D**, measured by PAM) and average Chl fluorescence lifetime (**E-F**, measured by TCSPC) were monitored in *npq4lhcsr1* mutant and its control strain (red and blue curves respectively) during 40 minutes of light fluctuations with periods of 1, 2, 4 and 10 minutes (**A/E**, **B/F**, **C/G** and **D/H** respectively) as described in **Fig. 1** of the main text. The solid lines represent the average of three biological replicates. Shaded regions represent standard deviation. For TCSPC data, each biological replicate was averaged from three technical replicates.

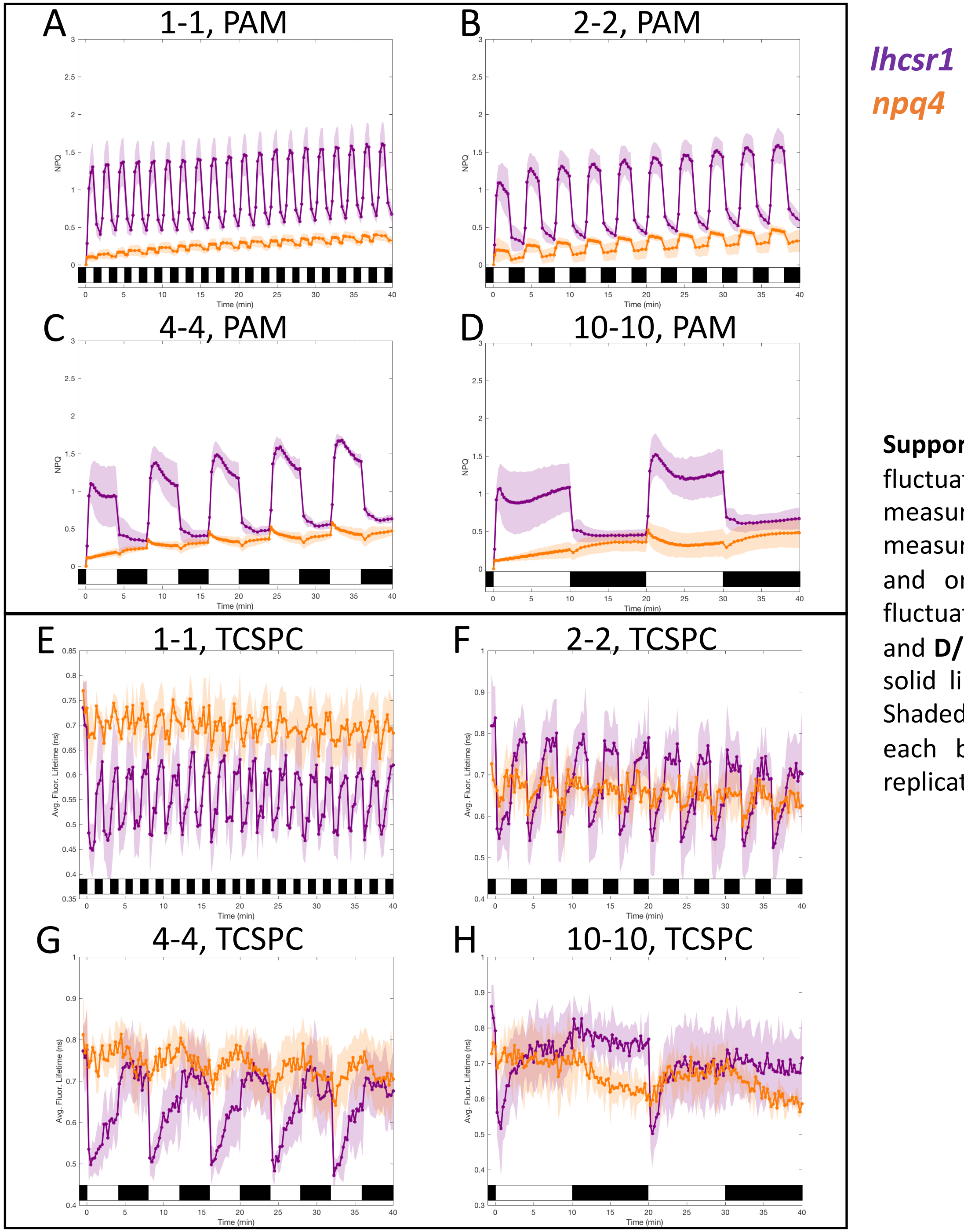


**Supporting Figure 14.** Quenching trajectories during light fluctuations in *lhcsr1* and *npq4*. The response of NPQ (**A-D**, measured by PAM) and average Chl fluorescence lifetime (**E-F**, measured by TCSPC) were monitored in *lhcsr1* and *npq4* (purple and orange curves respectively) during 40 minutes of light fluctuations with periods of 1, 2, 4 and 10 minutes (**A/E**, **B/F**, **C/G** and **D/H** respectively) as described in **Fig. 1** of the main text. The solid lines represent the average of three biological replicates. Shaded regions represent standard deviation. For TCSPC data, each biological replicate was averaged from three technical replicates.

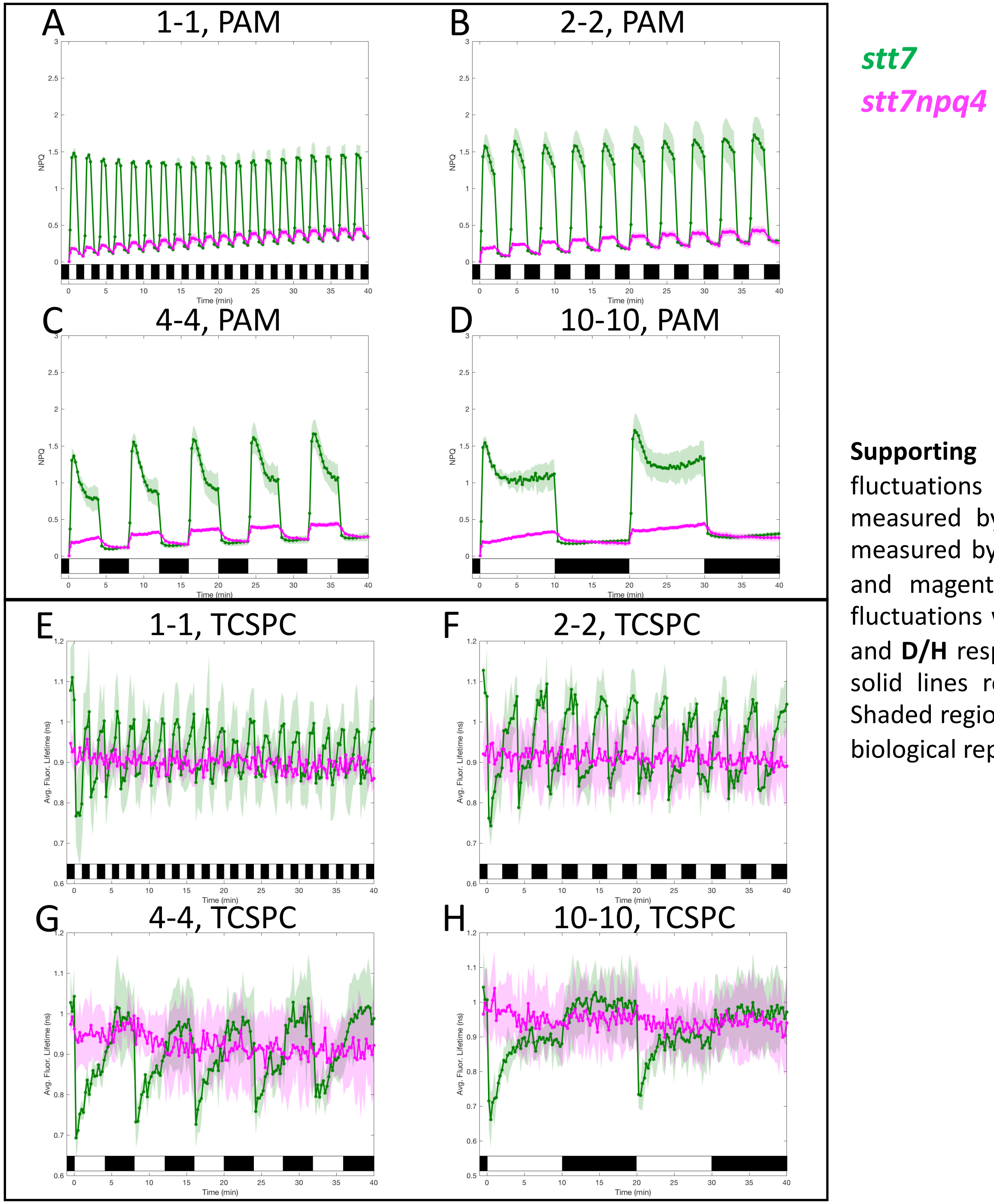


**Supporting Figure 15.** Quenching trajectories during light fluctuations in *stt7* and *stt7npq4*. The response of NPQ (**A-D**, measured by PAM) and average Chl fluorescence lifetime (**E-F**, measured by TCSPC) were monitored in *stt7* and *stt7npq4* (green and magenta curves respectively) during 40 minutes of light fluctuations with periods of 1, 2, 4 and 10 minutes (**A/E**, **B/F**, **C/G** and **D/H** respectively) as described in **Fig. 1** of the main text. The solid lines represent the average of three biological replicates. Shaded regions represent standard deviation. For TCSPC data, each biological replicate was averaged from three technical replicates.
